# Genetic variability in pathways associates with pesticide-induced nervous system disease in the United States

**DOI:** 10.1101/2023.09.25.559342

**Authors:** Marissa B. Kosnik, Philipp Antczak, Peter Fantke

**Author notes:** To whom correspondence should be addressed. Author e-mail addresses: Marissa B. Kosnik, Philipp Antczak Peter Fantke. Now at Department of Environmental Toxicology, Swiss Federal Institute of Aquatic Science and Technology, Eawag, 8600 Dübendorf, Switzerland (Marissa B. Kosnik).

## Abstract

Nervous system disease development following pesticide exposure can vary in a population, but genetic susceptibility to chemicals is poorly characterized. We developed a framework to build Chemical – SNP (single nucleotide polymorphism) – Disease linkages via biological pathways. We integrated these linkages with spatialized pesticide application data for the United States from 1992 – 2018 and nervous system disease rates for 2018 to characterize genetic variability in pesticide-induced nervous system disease. We found that the number of SNPs implicated per pesticide in US states positively correlates with disease incidence and prevalence for Alzheimer’s disease, Parkinson disease, and multiple sclerosis. Further, only 2% of pesticide sets used together over time overlapped between high disease occurrence and low disease occurrence states, with more SNPs implicated in pathways in high disease occurrence states. This supports that pesticides contribute to nervous system disease, and we developed priority lists of SNPs, pesticides, and pathways for further study.

## 1. Introduction

Interactions between environmental factors and genetics underlie the majority of chronic human diseases ^1^, and genome-wide association studies (GWAS) have identified thousands of loci that may describe inter-individual variability in disease progression. While some studies have described modifying effects of environmental factors like diet, alcohol, and smoking on disease outcomes, chemical exposures are rarely assessed in GWAS. This is in part due to the challenge of characterizing the spectrum of exposures an individual encounters (i.e. the exposome) and linking individual chemical exposures to particular genetic variants and disease outcomes ^2^. Because linking single nucleotide polymorphisms (SNPs) to specific environmental exposures often requires highly-exposed populations, pharmaceuticals are some of the only compounds this has been done for ^3^. Therefore, identifying genetic variants that may play a role in differential disease development following exposure to a wider range of chemicals is challenging.

Of the many chemicals humans may be exposed to, pesticides have a unique set of characteristics. Pesticides are ubiquitously used, toxic by design, and intentionally applied to agricultural crops and released into the environment ^4^. Occupational exposure to pesticides among agricultural workers is well-recognized, and high levels of pesticide residues have been found in drinking water and on food products available for consumption ^5^. Further, many different classes of pesticides have been implicated in nervous system diseases like Alzheimer’s disease, multiple sclerosis, and Parkinson disease through epidemiological studies of populations with chronic pesticide exposure ^6,7^. However, development of nervous system diseases following pesticide exposure is variable, suggesting some individuals may have more genetic susceptibility to pesticide-induced neurotoxicity ^8,9^. Given that pesticide use is expected to increase with population growth and climate change ^10^, it is important to determine the potential role of pesticides in differential development of nervous system disease to protect genetically susceptible populations.

The need to consider differential population susceptibility to chemical exposures is recognized in chemical risk assessment ^11,12^. Conventional methods relying on arbitrary safety factors (e.g., dividing a toxic dose by 10) cannot capture the complex role of human genetics in response to chemical exposures ^13^, and new approach methodologies (NAMs) have been proposed as a potential way to address this gap ^14^. We previously developed a dataset of Chemical-SNP-Disease linkages by integrating disparate datasets together ^15^. Using these novel linkages, we characterized differences in SNPs implicated by consumer products and distinguished potential mechanisms of toxicity for different disease classes ^16^. While this dataset helps characterize potential gene-environment interactions, the linkages were limited to SNPs located in genes and did not consider toxicity pathways at the Chemical-Disease intersections in forming a linkage. Here, we develop a new framework to form Chemical-SNP-Disease linkages using a toxicity pathway-based approach to identify SNPs implicated in differential population susceptibility to chemicals. Then, as an illustrative example of the value of this framework, we spatialize our newly-developed Pesticide-SNP-Disease linkages using pesticide application data and nervous system disease data for the United States (US). We focus on the US because it has publicly available spatialized pesticide application data, but this analysis can be repeated for other locations and chemical classes. Through our case study, we characterize the relationship between differential population susceptibility to pesticides and nervous system disease in the US. We then characterize the pesticides used and SNPs and genes implicated in different regions of the US and build priority lists of genetic variants, genes, disease pathways, and pesticides for further study of differential population susceptibility to pesticide-induced nervous system disease. This approach to form Chemical-SNP-Disease linkages—and the resulting Pesticide-SNP-Nervous system disease dataset—can serve as a starting point to characterize how chemical exposures lead to varied health outcomes in a population to incorporate inter-individual variability into chemical risk and impact assessment. Further, we demonstrate the applicability of this approach to explain the relationship between chemical exposure and disease in different geographic regions so that genetically susceptible populations can be better protected.

## 2. Methods

A general overview of the data integration process to form Chemical-Pathway-Gene-SNP-Disease linkages is shown in Figure 1. Briefly, Chemical-Gene data were integrated from high-throughput screening data and a literature-curated database. Pathway-Gene-Disease linkages were formed using gene enrichment. Chemical-Gene-Pathway-Disease linkages were formed based on Gene overlap in the Chemical-Gene and Pathway-Gene-Disease intersection using the Fisher’s exact test. SNPs were identified in Gene-Disease linkages. Additional SNPs located within 10000 bases of a SNP identified in a Gene-Disease linkage were also included. Chemicals were reduced to pesticides applied in the US, and diseases were reduced to six nervous system disease with incidence and prevalence data for the US. Linkages were spatialized based on where pesticides were applied in the US.

**Figure 1.**
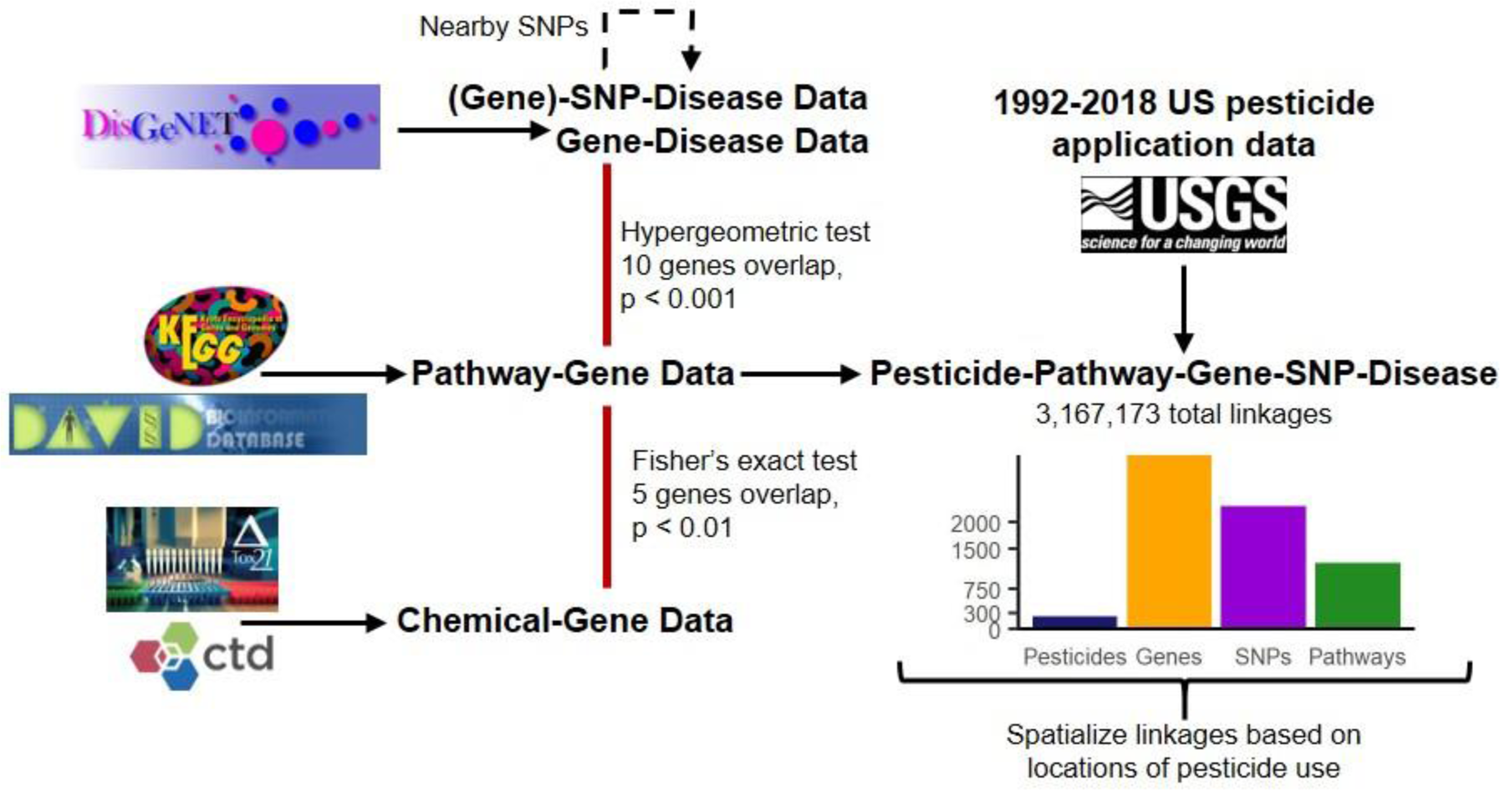
Overview of the Pesticide-Pathway-Gene-SNP-Disease linkage formation process. Data sources were integrated (left side of figure) to form linkages (right side of figure). Bar plots give the overall number of different data values forming the 3,167,173 Pesticide-Pathway-Gene-SNP-Disease linkages (Note: these linkages include cases with no SNPs implicated in a gene). The six nervous system diseases included are: Alzheimer’s disease, multiple sclerosis, Parkinson disease, brain neoplasms, epilepsy, and migraine disorders.

### 2.1. Chemical-gene data integration

High-throughput screening (HTS) chemical-gene data from Tox21/ToxCast ^17^ and literature-curated data from Comparative Toxicogenomics Database (CTD) ^18^ were merged to increase the chemical-gene associations used in the analysis, following the general protocol outlined in Kosnik et al. 2019 ^19^. Details of this approach are in Section S1. In brief, HTS and CTD chemical-gene associations were merged using CAS Registry Numbers (CASRN) and Entrez Gene IDs. The integrated HTS/CTD dataset was then reduced to pesticides used in the US as reported in the United States Geologic Survey (https://water.usgs.gov/nawqa/pnsp/usage), which provides the amount of pesticide applied in kilograms per county in the US in 1992 – 2018. The integrated dataset had 21,401 unique chemical-gene associations describing 338 chemicals and 12,942 genes.

### 2.2. Pathway-Gene-Disease integration

Pathway disease linkages were formed through overrepresentation analysis (ORA) based on the gene space in the pathway-disease intersection. Details on the integration and sensitivity analysis are described in Section S2. Briefly, Pathway-Gene data were collected (e.g., from DAVID Knowledgebase ^20^), and integrated with Gene-Disease data from DisGeNET ^21^ to form enriched Pathway-Gene-Disease linkages. Of the 903 nervous system diseases in the dataset (Medical Subject Headings class C10), six with disease matches in the Global Burden of Disease (GBD) were included for further analysis: Alzheimer’s disease, multiple sclerosis, Parkinson disease, brain neoplasms, epilepsy, and migraine disorders.

### 2.3. Chemical-Gene-Pathway-Disease integration

Chemical-Disease linkages were formed based on a significant subset of genes present in the Chemical-Gene and Pathway-Gene-Disease datasets, suggesting that a chemical could act through the given pathway to lead to a disease outcome. Significant Chemical-Disease linkages were formed using Fisher’s exact test. Details on the data integration and sensitivity analysis are in Section S3. The final set of Chemical-Pathway-Gene-Disease linkages (or Chemical-Disease linkages for simplicity) were formed by requiring a Pathway-Disease linkage to be enriched with an adjusted p-value < 0.001 and 10 genes overlap and Chemical-Pathway-Disease linkages with a p-value < 0.01 and 5 genes overlapping.

### 2.4. Chemical-SNP-Disease linkages

SNPs were considered implicated in a Chemical-Disease linkage if they were implicated in the same disease and in or near a gene implicated by both the chemical and the disease. SNP-Disease data were collected using the disgenet2r package version 0.99.3 ^21^. For each Chemical-Disease linkage, all genes implicated at the intersection were identified (based on pathways as described in Section 5.3). Then, all SNPs located in the same genes and implicated in the same diseases were identified. This yielded Chemical-Pathway-Gene-SNP-Disease linkages (or Chemical-Gene-Disease/Chemical-SNP-Disease linkages depending on which data are being considered in the linkage intersection).

Then, to identify intergenic SNPs, SNPs located within 10000 bases of another SNP implicated in the same disease (the general maximum value accepted as a SNP association for a given gene ^22^) and implicated in a Chemical-Pathway-Gene-Disease linkage were identified.

These SNPs were considered potentially implicated in the same Chemical-Disease association as the original (located in a gene) SNP. Variant consequences were available in DisGeNET, and consequence severity was determined from Ensembl (https://www.ensembl.org/info/genome/variation/prediction/predicted_data.html). Chemical-Gene-Disease linkages without SNPs were also kept to identify genes that are important in disease outcomes or may have unknown, disease-relevant SNPs in a population. The full set of Pesticide-Pathway-Gene-SNP-Disease linkages is available in Table S2. The previous, non-pathway-based integration of the dataset included multiple evaluation steps between external SNP-Disease and Chemical-SNP-Disease linkages ^15,16^. Additional evaluation against external Pesticide-SNP-Disease linkages is shown in Table S1. This increases confidence in the linkages formed between pesticides, SNPs, and diseases in the present analysis.

### 2.5. Spatial assessment of Pesticide-SNP-Disease linkages

Pesticide-Gene-SNP-Disease linkages were mapped to different US states based on where the pesticide was reported to be used in the USGS dataset (https://water.usgs.gov/nawqa/pnsp/usage/maps/). Disease data for the US was collected from the Global Burden of Disease for 1992 – 2018 (https://www.healthdata.org/gbd/2019), the same years as pesticide application data from USGS. Disease incidence and prevalence rates were collected for “Alzheimer’s disease and other dementias”, “Parkinson’s disease”, “Idiopathic epilepsy”, “Multiple sclerosis”, “Brain and central nervous system cancer “, and “Migraine” and matched to the corresponding diseases in the Pesticide-SNP-Disease dataset: “Alzheimer’s Disease”, “Parkinson Disease”, “Epilepsy”, “Multiple Sclerosis”, “Brain Neoplasms”, and “Migraine Disorders.”

## 3. Results

We generated Pesticide-Pathway-Gene-SNP-Disease linkages by integrating multiple publicly-available sources. For simplicity these are termed Pesticide-SNP-Disease linkages or Pesticide-Gene-Disease linkages if no SNP was implicated. The final dataset includes 234 pesticides implicating 2304 total SNPs and 3244 total genes across 1230 toxicity pathways in six nervous system diseases of interest: Alzheimer’s disease, Parkinson disease, multiple sclerosis, brain neoplasms, epilepsy, and migraine disorders (Figure 1). While there are limited data on SNP-Disease or Gene-Disease interactions in pesticide-exposed populations, we found good agreement between the linkages we formed and SNPs implicated in differential population susceptibility to pesticides in the literature (Table S1). For example, multiple studies identified SNPs in HLA-DRA, ABCB1, APEX, PON1, and PPARGC1A as influencing pesticide-induced Parkinson Disease in a population, and our approach identified these same SNPs in the Pesticide-Disease intersection^23–30^. The full set of Pesticide-Pathway-Gene-SNP-Disease linkages is available in Table S2 and an overview of these linkages is provided in Section S4.

### 3.1. Differential population susceptibility to pesticide exposures may influence nervous system disease rates in the US

To characterize the potential role of genetic susceptibility in pesticide-induced nervous system diseases, we used spatially-refined pesticide application data for the contiguous US from 1992 – 2018 to map the Pesticide-SNP-Disease linkages to different states based on where each pesticide was applied (Section S5). To assess the cumulative effect of pesticide exposure on SNPs/genes in a disease outcome, each instance of a pesticide acting on a SNP/gene was considered a “hit.” Because multiple pesticides can implicate the same SNP/gene, and pesticides can be applied in multiple years, this means the same SNP/gene may be implicated multiple times in the same region (see Section S5.1 for details). This enabled us to identify US states where genetically susceptible populations may be more vulnerable to pesticide exposures. Using these spatialized Pesticide-SNP linkages, we looked at the relationship between pesticide use, implicated genes/SNPs, and disease outcomes in the United States. The rate of disease incidence and prevalence by state for the six diseases under study was collected from the Global Burden of Disease for the years 1992 – 2018. To include as many years of pesticide application as possible representing chronic pesticide exposure preceding a disease, we relate pesticide use from 1992 – 2018 to disease rates per 100000 people for the year 2018 for the remainder of the analysis and assume state-to-state migration does not impact the analysis (see discussion of other years of disease data and migration rates in Section S6-S7).

To assess if pesticide use in a state may influence disease occurrence, we compared the amount of pesticide applied per state (kilograms/square mile of the total state area) to disease incidence and prevalence rates for 2018, and found no clear association (Figure S7). This is not surprising since the amount of pesticide applied does not necessarily explain potential biological effects and regulatory agencies strive to minimize exposure of humans to these compounds.

Interestingly however, by relating the number of SNP hits implicated per pesticide per year for 1992 – 2018 to disease occurrence in 2018, we found a significant, positive correlation for Alzheimer’s disease, multiple sclerosis, and Parkinson disease (spearman’s rho > 0.3, correlation p < 0.05, Fig 2, see Figure S4 for how SNP hits per pesticide per year are calculated, Figure S8 for all disease correlations). We also see a similar relationship between disease incidence/prevalence and the number of gene hits per pesticide applied per year, the average number of SNP hits per pesticide across years, and the number of unique SNPs (not hits) per pesticide (spearman’s rho > 0.3, correlation p < 0.05, Figure S9-S10). When evaluating whether just the number of pesticide applications or the total number of SNP hits implicated in a state shows a similar correlation with disease incidence/prevalence we found that these resulted in negative correlations (Figures S14-S15). This suggests that, as expected, it is not the raw number of times pesticides are applied nor just the number of SNPs that are being hit that explains increased disease incidence/prevalence but rather the combination of SNPs and the pesticides that associate with them. This is in line with the concept that the “dose makes the poison” which we hypothesize also applies to SNPs in a population wherein the same pesticide frequently associated with the same SNP has a greater effect than many pesticides acting on many different SNPs. Based on the positive correlations we identified, pesticides applied in states with higher nervous system disease incidence/prevalence implicate more unique SNPs per pesticide on average and pesticides may act on these SNPs more frequently. To account for SNPs affected by pesticide application over time, we focus on SNP hits per pesticide per year for the remainder of the analysis (Figure 2A).

**Figure 2.**
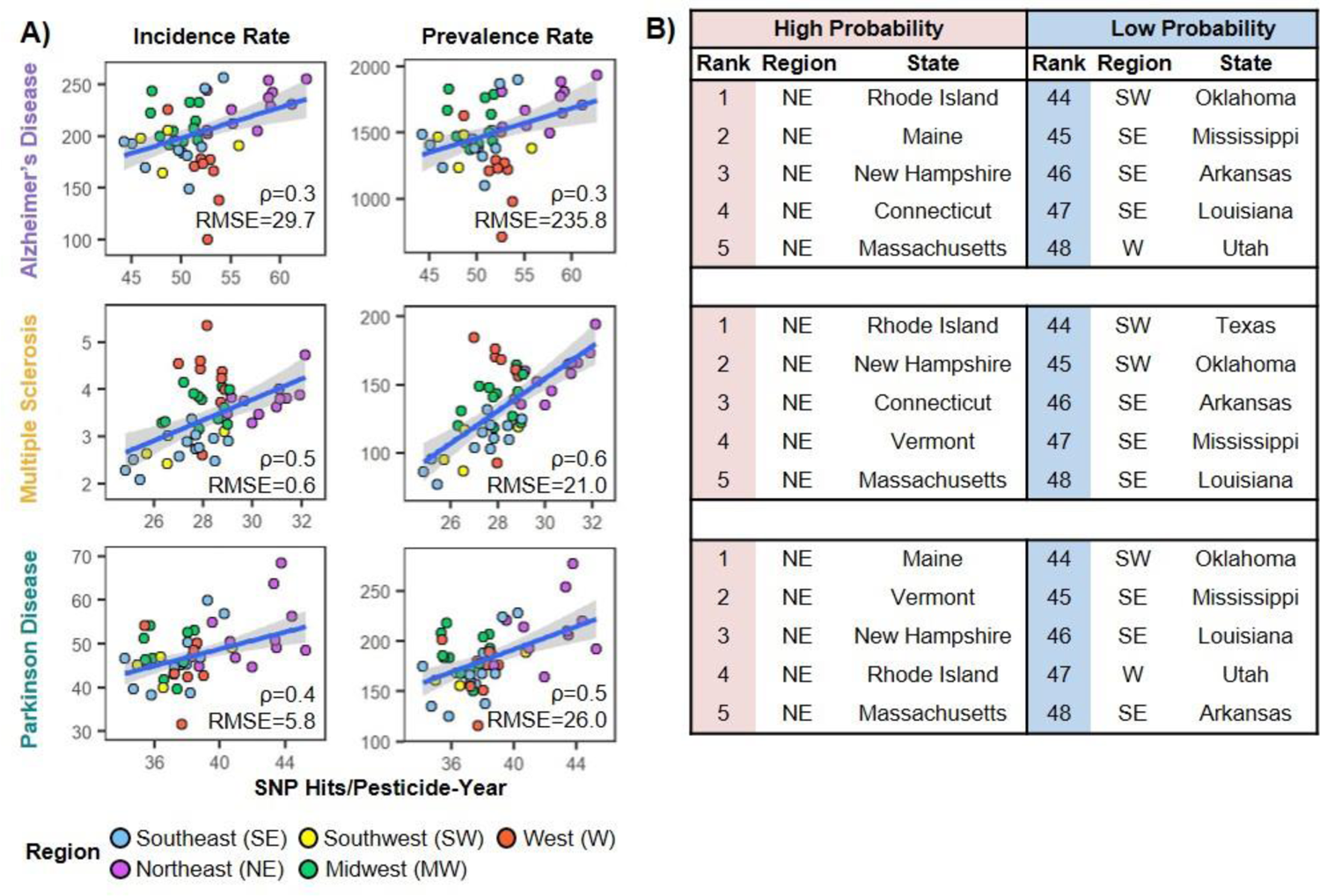
States using pesticides with more SNPs implicated have higher disease incidence and prevalence. A) Scatterplots of SNP hits/pesticide-year for pesticides applied in 1992 – 2018 vs disease incidence and prevalence in 2018. Each point is a US state. Colors = geographic region of each state. Trend line = robust linear model, RMSE = root mean square error, ρ = Spearman’s rank correlation coefficient. All correlations are significant p < 0.05. B) High and low probability states based on the number of SNP hits per pesticide-year and disease incidence/prevalence.

Interestingly, a bootstrapping approach highlighted that multiple sclerosis and Parkinson’s disease correlation coefficients were significantly higher than expected by chance (see Section S8 for more details). In addition, the MAE and RMSE were significantly lower for Alzheimer’s disease, multiple sclerosis, and Parkinson disease suggesting that in these diseases pesticide usage may play a significant role in disease incidence and prevalence.

### 3.2. Multiple pesticides with variable SNP linkages may contribute to disease occurrence in each state

The observation that SNP hits per pesticide for Alzheimer’s disease, Parkinson’s disease, and multiple sclerosis resulted in a positive correlation led us to hypothesize that the relationship between the SNPs and pesticides is more important in disease occurrence than just the types of pesticides or SNPs implicated per state. To assess whether individual pesticides were driving the associations between disease occurrence and SNP hits per pesticide, we applied a regularized regression approach (LASSO – Section S9 for details) to identify which predictors may contribute more significantly to the association. For Alzheimer’s disease (218/227), multiple sclerosis (205/216), and Parkinson’s disease (212/223) most pesticides were selected at least 70 times out of 100 LASSO runs (full set of coefficients in Table S3).

For Alzheimer’s disease, the herbicide alachlor, insecticide clothianidin, fungicides thiophanate and azoxystrobin, and hydrogen peroxide received the largest coefficients (0.65-0.81). The top five pesticides for multiple sclerosis were insecticides ethoprop, deltamethrin, acetamiprid, and tebufenozide, and fungicide zoxamide (0.68-0.88) and Parkinson disease had herbicides alachlor, tribenuron-methyl, and oxyfluorfen, fungicide thiophanate, and hydrogen peroxide as the top five pesticides (0.61-0.77). While the ranks of the top 100 pesticide coefficients differed between the three diseases (Mann-Whitney test p < 0.05), the ranks of all pesticide coefficients did not significantly differ. This suggests some differences in pesticides driving the correlation between disease occurrence and Pesticide-SNP hits in US states. However, this analysis cannot disentangle the exact relationship between pesticide use and high disease occurrence.

### 3.3. Different Pesticide-Gene-SNP-Disease linkages are implicated over time in high and low probability regions

To determine what may underlie the relationship between disease occurrence and Pesticide-SNP linkages implicated in differential population response, we analyzed the top five and bottom five states with the highest and lowest values for disease incidence/prevalence and SNP hits per pesticide-year, i.e. those closest to the top right and bottom left corners of the disease correlation plots (Figure 2A-B, referred to as high and low probability states respectively) for the three correlated diseases: Alzheimer’s disease, multiple sclerosis, and Parkinson disease (Section S10 for details).

Across the three diseases, Rhode Island, New Hampshire, and Massachusetts were high probability states for all three diseases, and Oklahoma, Mississippi, Arkansas, and Louisiana were low probability states for all three diseases. Some pesticides were applied in high probability states between 1992 – 2018, but were never used in low probability states, including chlorpropham, flumetralin, and 2,6-dichlorobenzonitrile which were implicated in all three nervous system diseases through our linkage formation process. However, despite the unique pesticides used in high probability states, no SNPs were implicated in high probability states that were not also implicated by pesticides used in low probability states. Therefore, we hypothesize that, even if the same SNPs are implicated by pesticides in high and low probability states, the frequency and efficacy at which pesticides act on SNPs differs. For example, the fungicides zoxamide and triflumizole, and insecticide pyridaben were applied in at least eight years in all high probability states for all three diseases, but were applied in only one of the low probability states for each disease (Utah for Alzheimer’s disease and Parkinson disease, Texas for multiple sclerosis). This means the SNPs and genes implicated by these pesticides would be more frequent hits in high probability states than low probability states. We therefore hypothesize that pesticide use over time (and thus, SNPs/genes hit over time) may underlie the differences observed in disease occurrence between high and low probability states.

### 3.4. Different sets of pesticides were used together over time in high-probability and low-probability regions

To differentiate pesticide use patterns between the high and low probability states, we performed frequent itemset mining on the annual USGS data per state. For this process, each individual year of pesticide application data (1992 – 2018) was treated as a unique transaction with all the pesticides applied in a state in that year considered input items (Figure S20). Association rule directionality was not considered. This analysis identified sets of 2 – 4 pesticides that were applied together in at least five years in the same state (i.e. pesticide usage patterns – See Section S11 for details).

Across all 48 states included in the analysis, 1,341,845 unique rules (i.e. sets of 2 – 4 pesticides applied together) were identified with frequently applied pesticides like glyphosate, atrazine, and 2,4,D implicated in multiple rules across states. However, other pesticides were used differently between states (e.g. zoxamide) leading to different association rules. Across diseases, the high probability states had 72,588 total rules while the low probability states had 250,670 total rules suggesting a more variable pesticide usage (Figure 3A). Comparing association rules in the high and low probability states, < 2% of association rules were the same, meaning the pesticides most frequently used together in 1992 – 2018 differ between the states (hypergeometric test p = 1, meaning no significant overlap between rules in high and low probability states). Comparing the modes of action (MoAs) of pesticides in these association rules, insecticides were used most frequently in high probability state rules while herbicides were most frequently implicated in low probability state rules (Figure 3A).

**Figure 3.**
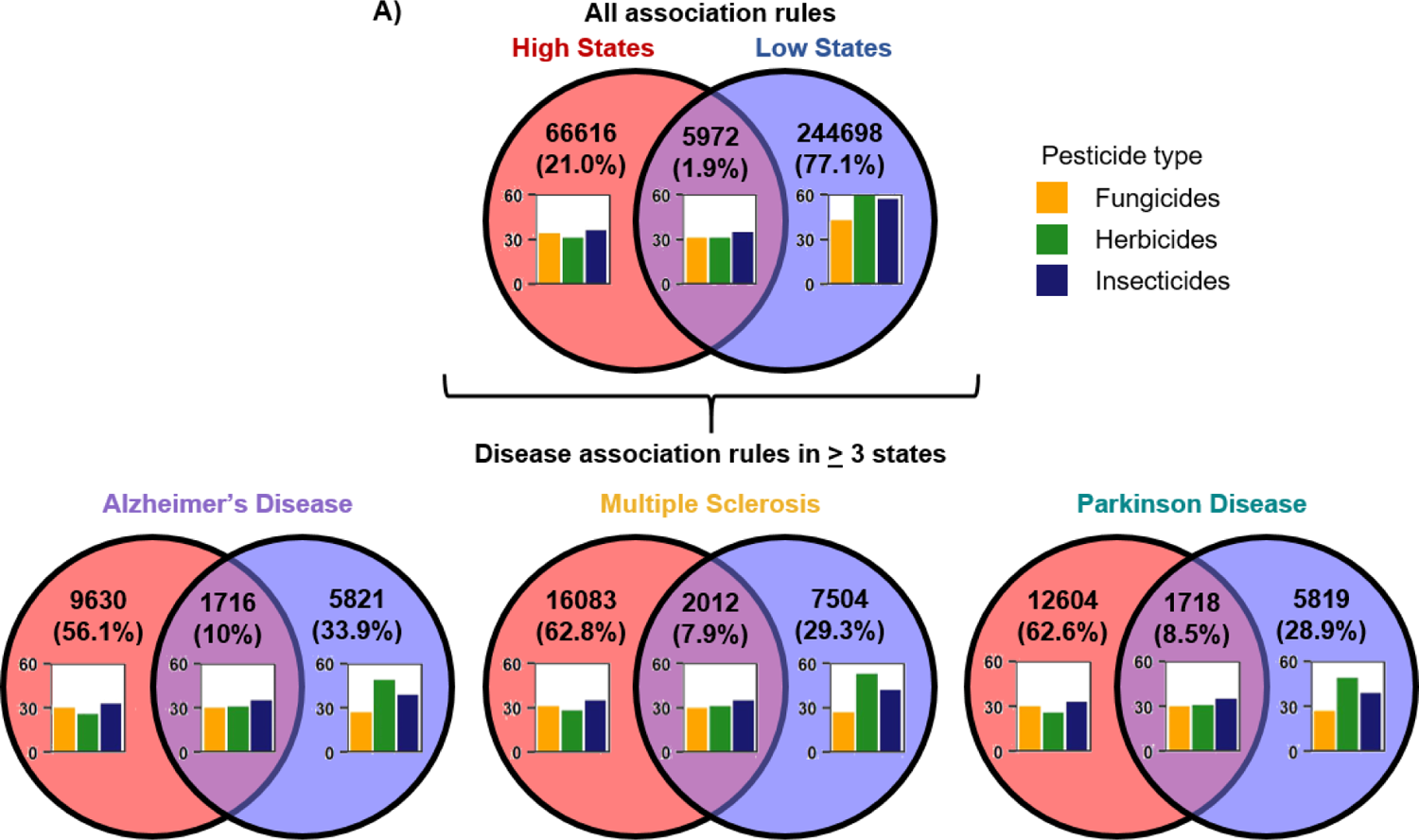
Different pesticide combinations were used in high probability and low probability states in 1992 – 2018. A) Venn diagram of all association rules in high or low probability states or both showing pesticide use patterns in 1992 – 2018. B) Venn diagrams of disease-specific association rules present in three or more states. The pesticide types for pesticides in the association rules are in each bar chart (Note: pesticide type not available for all pesticides).

To characterize differences in pesticide use in high and low probability states for each disease, we mapped the identified association rules for each state to Alzheimer’s disease, multiple sclerosis, and Parkinson disease based on the pesticides implicated in each disease in our dataset (Section S11). Then, we reduced the association rules to those present in at least three states to ensure the rule is characteristic of high or low probability states rather than just representing a pesticide use pattern in one state (Figure 3B). As with the overall set of association rules across diseases, the association rules were significantly different between high and low probability states for each disease (hypergeometric test p = 1), and insecticides were implicated most in high probability states while herbicides were implicated most in low probability states. Interestingly, by requiring at least three states to have an association rule in common, the number of association rules in low probability states dropped below the number of association rules in high probability states for each disease (SI Figure S21). This further suggests that the pesticide use patterns in the high probability states may explain the relationship we see between pesticides, SNPs, and nervous system disease rates compared to the variable pesticide use patterns in low probability states. The full set of association rules in high probability states is in Table S4.

### 3.5. More SNPs and genes are frequently implicated in disease pathways for high probability states than low probability states

To prioritize SNPs, genes, and pathways that may drive differences in the relationship between the Pesticide-SNP hits and disease occurrence in high and low probability states, we identified the SNPs and genes that were more frequent hits in high probability states than low probability states (i.e. those SNPs and genes that pesticides acted on more in high probability states) and the corresponding disease pathways. Based on the increased disease incidence/prevalence in high probability states, these SNPs, genes, and pathways may be more important in disease outcomes. We found 1386 SNPs and 2129 genes that were more frequent hits in high probability states than low probability states compared to 402 SNPs and 475 genes that were more frequent hits in low probability states than high probability states. This supports our hypothesis that variable pesticide use in low probability states results in variable SNP/gene hits compared to high probability states where the same SNPs and genes are frequently implicated. We ranked the high priority SNPs and genes based on the hit frequency and the number of association rules implicating each SNP and gene (Section S12 for details). The full set of high priority SNPs and genes is in Table S5.

While some SNPs overlapped between the three diseases, the majority of the priority SNPs were unique to each disease (Figure 4A). Of the 1386 priority SNPs, 82 were found in two overlapping diseases (mostly between Alzheimer’s disease and Parkinson disease), and only 14 were in all three diseases with rs1800795 in IL6 in the top 25 SNPs for all three diseases. The ten highest priority SNPs for Alzheimer’s disease include several SNPs in CYP19A1, and one in MAPK1 and CD69. Multiple sclerosis also has a high priority SNP in MAPK1, two in CD69, and several in CYP24A1 and IL1A/B while Parkinson disease had two in SIRT1 and IL1B, and one in HMOX1, IL6, and AKT1. While most of the high priority SNPs are modifiers according to Ensembl classifications, the top SNPs for all three diseases are moderate severity, and three of the top 25 SNPs for Parkinson disease are high severity (rs1485215606 in CASP3, rs3832852 in A2M, and rs1470543813 in IGFALS).

**Figure 4.**
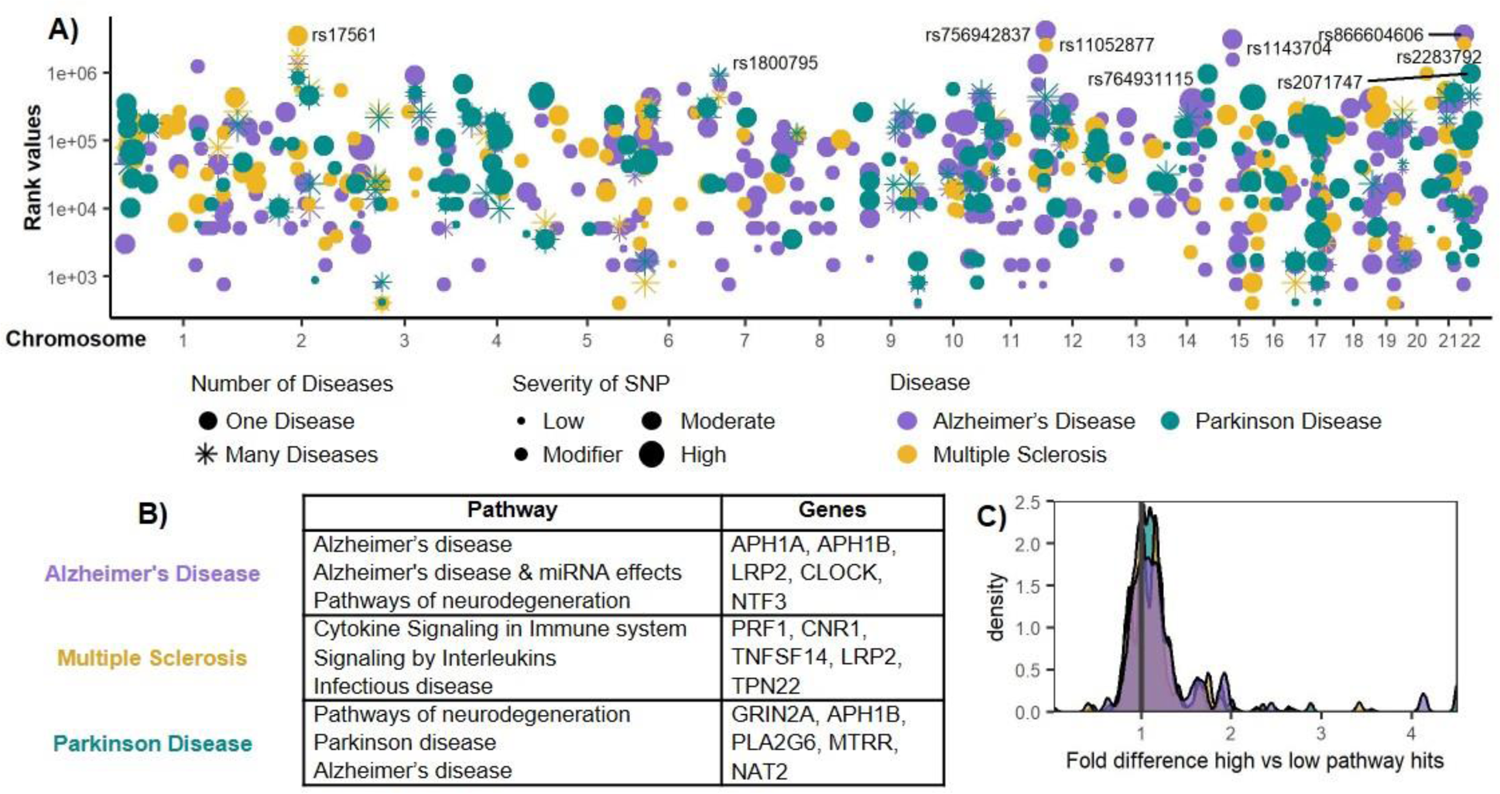
More SNPs and genes are frequent hits of pesticides applied in high probability states than low probability states. A) Manhattan plot of priority SNPs more frequently implicated in high probability states than low probability states. Y-axis = SNP rank value determined from the number of SNP hits in 1992 – 2018 and the number of association rules with pesticides implicating that SNP. Circles = SNPs only present in one disease, Stars = SNPs present in multiple diseases. SNP size = severity of the variant (e.g., stop gained variant). B) Table of pathways and genes with the greatest fold hit differences in high probability states compared to low probability states. Note: the genes may not all be present in the given pathways. C) Fold greater (> 1) or lower (< 1) gene hits in a disease pathway in high probability states compared to lower probability states. The color legend is the same as in A.

To determine the pathways that may underlie the increased disease occurrence in high probability states, pathways were ranked by the fold difference in SNP/gene hits between high probability states versus low probability states. For Alzheimer’s disease, the top-ranked pathways for the high probability states include Alzheimer’s disease and pathways of neurodegeneration while multiple sclerosis had immune system-related pathways like signaling by interleukins and Parkinson disease pathways include Parkinson disease and pathways of neurodegeneration (Figure 4B). The full set of ranked pathways implicated in high probability states is in Table S6. Of the 1081 pathways implicated by the 1386 priority SNPs and 2129 genes for high probability states, only 21 pathways were not also implicated in low probability states.

Therefore, we suspect that the SNPs/genes implicated in the high and low probability states lead to different activity of the same pathways following exposure to a pesticide. Figure 3C shows the fold greater or lower Gene-Pathway hits from pesticides used in high probability or low probability states (i.e. how many times more or less a gene was implicated by pesticides in a disease pathway in high states versus low states). For both Alzheimer’s disease and Parkinson disease, 80% of pathways had more SNP/gene hits in high probability states than low probability states while 67% of multiple sclerosis pathways had more SNP/gene hits in high probability states than low probability states. This may mean there is more potential gene/SNP activity in disease pathways from pesticides used in high probability states compared to low probability states.

### 3.6. Many pesticides implicate high-priority SNPs and are applied in greater amounts in high probability states than low probability states

To prioritize pesticides that may drive the differences between high and low probability states, we identified pesticides applied in greater quantities in high probability states than low probability states and ranked these by the mass applied, number of implicated high priority SNPs/genes (SNPs in Figure 4A), and presence in association rules, meaning they were applied in at least five years in 1992 – 2018 (see Section S12 for details). This resulted in 40 pesticides of interest (Figure 5, Table S7).

**Figure 5.**
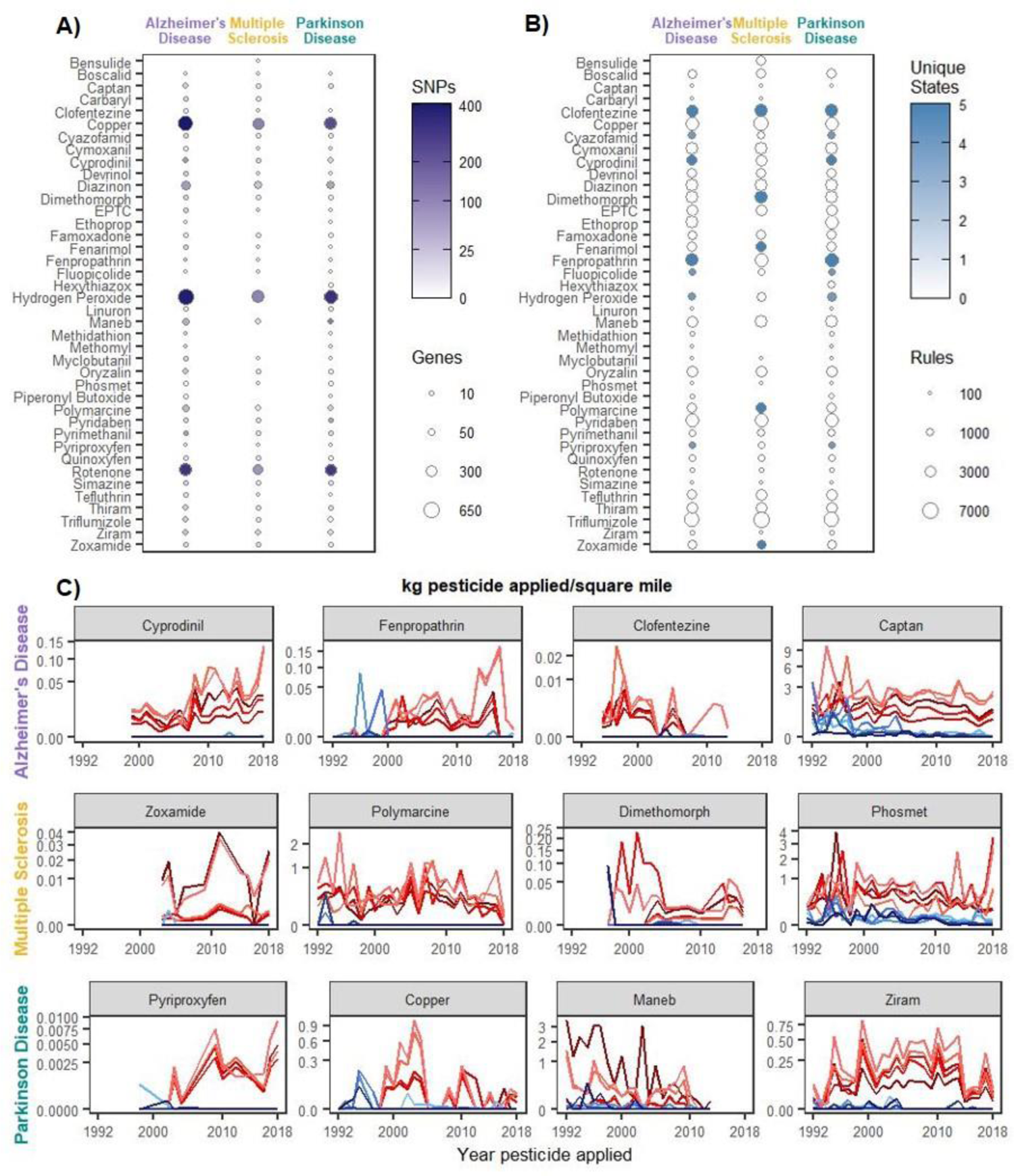
Priority pesticide linkages and application patterns over time. A) The number of priority genes and SNPs (Table S5) in priority Pesticide-Disease linkages. B) The number of association rules (Table S4) involving priority pesticides. Unique states refer to the number of states with a pesticide used in rules that were uniquely found in high probability states and no low probability states. C) Sample priority pesticide application in kg/square mile for 1992 – 2018. Red trend lines = high probability states, blue trend lines = low probability states.

The insecticide clofentezine was in the top three prioritized pesticide for all three diseases and uniquely implicated in association rules in all high probability states. While clofentezine was used in some low probability states, it was applied in three or fewer years compared to 11 – 14 years in high probability states. Additionally, the fungicide cyprodinil and insecticide fenpropathrin were in the top three pesticides for both Parkinson disease and Alzheimer’s disease and were uniquely implicated in association rules for all five high probability states for both diseases. Copper, hydrogen peroxide, and rotenone were among the top ten highest priority pesticides for all three diseases, with polymarcine (metiram), cyprodinil, diazinon, maneb, and ziram, among the top 15 high priority pesticides for all three diseases. Based on potential to cause harm in humans from common targets for a given pesticide MoA, we would expect more insecticides and/or fungicides to be prioritized in high probability states than low probability states. Compared to the full set of pesticides used across all states, the priority list for Alzheimer’s disease was enriched for insecticides with acetylcholinesterase inhibitor MoAs, and all three diseases were enriched for fungicides targeting amino acid protein synthesis, mitosis and cell division, and multi-site contact activity (hypergeometric test p < 0.05). Only six herbicides were identified with priority across all three diseases, and had no enriched MoAs compared to the full set of pesticides included in the analysis.

## 4. Discussion

We integrated diverse datasets to form Chemical-Pathway-Gene-SNP-Disease linkages to better account for individual human genetic susceptibility to chemical exposures in risk assessment. These linkages do not depend on existing evidence that a chemical leads to an adverse outcome in genetically susceptible individuals, which is important for predicting chemical effects before an adverse outcome occurs to protect potentially-exposed populations.

With the huge number of marketed chemicals and the concern of how chemical exposures contribute to adverse outcomes, NAMs are necessary to address the growing number of possible chemical-adverse outcome combinations and account for inter-individual variability ^14^. One proposed approach that differs from ours is to identify SNPs in Adverse Outcome Pathway (AOP) genes using data from the AOP Wiki ^31^. Romano et al. 2023 used this approach to identify SNPs of interest in liver cancer by linking UK Biobank genetic data to liver cancer AOPs and identifying SNPs implicated at the intersection ^32^. Future analyses could combine this AOP-based approach with our approach to increase confidence in the validity of associations formed using each respective method or even identify new AOPs from our set of Pesticide-Pathway-Gene-SNP-Disease linkages.

We found evidence that pesticides contribute to Alzheimer’s disease, multiple sclerosis, and Parkinson disease, and that the SNPs and genes implicated at the Pesticide-Disease intersection explain the extent to which this occurs. To our knowledge, our approach using spatialized pesticide application data to identify SNPs that may be relevant to disease occurrence in different geographic regions has not been done before. Eccles et al. 2023 used spatialized exposure data to predict the potential for different geographic regions to be affected by chemicals^33^. While the focus of their methods was to characterize exposure to chemical mixtures based on potential molecular target perturbations (without consideration of specific SNPs, specific chemicals, or specific adverse outcomes), integrating some of their methods with our spatialized Pesticide-SNP-Disease linkages represents one approach to quantify the potential for the pesticides we identified to trigger the implicated pathways in disease progression.

We developed priority lists of SNPs and genes more frequently implicated in high probability regions than low probability regions, pesticides that may be most likely to act on these SNPs/genes, and biological pathways that may be most important in driving disease occurrence. These lists can serve as a starting point to explore the role of pesticides in nervous system disease and to account for inter-individual variability in chemical risk assessment. Of the 1386 priority SNPs identified, very few SNPs overlapped between the diseases, suggesting that the points of differential population susceptibility to chemical exposures vary between diseases. This is in line with the different toxicity mechanisms of importance we identified between the diseases in our list of priority pathways. Many of the top SNPs and genes we identified have extensive external support for their importance in the identified diseases. For example, one of the highest-ranked gene for Alzheimer’s disease is MAPK1, which is important in Alzheimer’s disease progression ^34^ and other nervous system diseases ^35^. Further, with the exception of Alzheimer’s Disease, all SNPs/genes we found with literature supporting their role in pesticide-induced nervous system disease (Table S1, e.g., rs3129882 in HLA-DRA, ^23^) were identified as high priority SNPs/genes in our analysis.

We identified priority pesticides associated with nervous system diseases based on their potential to affect susceptible individuals. Rotenone was identified with priority in all three diseases, and is a well-recognized contributor to neurotoxicity, with a “rotenone model for Parkinson disease” being a common model to mimic environmental toxicant-induced Parkinson disease ^36,37^. We also identified cyprodinil as a high priority pesticide in all three diseases, and the top priority pesticide for Alzheimer’s disease and Parkinson disease. Cyprodinil applied in mixtures has been found to cause adverse nervous system effects in mice, which supports our hypothesis that the combination of pesticides applied in high priority states may be more important in the relationship between disease occurrence and SNP hits per pesticide-year than any individual pesticide ^38^. Additionally, combinations of pesticides frequently affect BACE1—a critical mediator in Alzheimer’s disease—that we identified as a high priority gene for further study ^38,39^. The pyrethroid fenpropathrin was also in the top three prioritized pesticides for Alzheimer’s disease and Parkinson disease, and has been found to induce loss of dopaminergic neurons and mimic some of the pathologic features of Parkinson disease in mice ^40^. Additionally, both Polymarcine (metiram) and maneb were prioritized for all three diseases. Metiram and maneb are both ethylenebisdithiocarbamate (EBDC) fungicides, and while polymarcine toxicity is understudied compared to other EBDCs, both rodent models ^41^ and epidemiological studies have implicated maneb in Parkinson disease, especially with co-exposure to paraquat (which was applied in all years in all states in our study) ^42,43^. We also identified metiram as a high priority chemical for multiple sclerosis and, while metiram has not been linked to multiple sclerosis in the literature, it has been linked to amyotrophic lateral sclerosis (ALS), so it may also be worth investigating further ^44^. Lastly, we identified copper among the top ten priority pesticides in all three diseases. Copper is an essential metal with a pivotal role in the human nervous system, and is used heavily in organic farming ^45^. However, there is some evidence that exposure to copper may increase the risk for Parkinson disease ^46,47^ and multiple studies have found environmental copper exposure increases the risk for Alzheimer’s disease in susceptible individuals ^48,49^. An epidemiological analysis found that individuals with CYP2D6 polymorphisms and the GSTP1 SNPs rs1695 and rs1138272 may be more susceptible to copper in the development of Alzheimer’s disease ^50^, and we also identified CYP2D6 and GSTP1 SNPs with high priority in copper-induced Alzheimer’s disease. This literature support combined with our findings highlights copper as an important compound for further investigation in differential development of nervous system disease.

We found that some of the compounds implicated in high-risk rules for each disease did not have a linkage with the disease in our starting Pesticide-Gene-SNP-Disease dataset. For example, all three diseases implicated vincozolin with high risk based on pesticide usage (no low probability states had vinclozolin in an association rule), but vinclozolin was not linked to these diseases in our dataset. Vinclozolin is an antiandrogenic fungicide that has been implicated in disruption of sexual differentiation and reproductive function ^51^. While there is no existing evidence that vinclozolin is linked to nervous system disease, it has been found to promote epigenetic transgenerational effects, which could influence individual differences in drug response ^52^, modify responses to other exposures, or influence disease inheritance ^51^. Therefore, future analyses should consider the role of vinclozolin—potentially in combination with other pesticides—in differential development of Alzheimer’s disease, multiple sclerosis, and Parkinson disease. Future analyses can also expand our work by analyzing more of the association rule differences (representing different pesticide use combinations) between high and low probability states.

The primary limitation in our analysis is the assumption that pesticides used in a geographic region affect individuals living in that location. Likely routes of exposure are not considered (e.g., oral versus inhalation), nor the likelihood that an individual would come in contact with a pesticide based on its physicochemical properties (e.g., persistence). Future analyses could focus on likely highly-exposed groups (e.g., farmers, bystanders, people who eat more locally-grown produce), incorporate trade data on crops, or incorporate likely pesticide minimum residue levels to better predict actual pesticide concentrations that individuals are exposed to. Additionally, it is unlikely that the diseases we are analyzing only occur due to pesticide exposure, so it is possible that some of the associations we found have other factors underlying them in our or other more epidemiological analyses (e.g., diet, lifestyle, other chemical exposures). While these are important considerations to determine how individuals in a region are exposed to pesticides, our preliminary approach is still capturing a signal between spatialized pesticide usage and disease occurrence which may indicate local exposure pathways are still relevant.

## 5. Conclusions

We formed a dataset of Pesticide-Pathway-Gene-SNP-Disease linkages to characterize the mechanisms by which pesticides can lead to nervous system disease in individuals with different genetic susceptibility. We found that pesticides may contribute to Alzheimer’s disease, multiple sclerosis, and Parkinson disease in the United States, and the priority lists of SNPs, genes, pathways, and pesticides we developed identify potentially important contributors to this relationship. This work represents a novel approach to consider inter-individual variability in disease outcomes where existing Chemical-SNP-Disease associations are limited. The approach we demonstrate for data integration and subsequent analysis of Chemical-Pathway-Gene-SNP-Disease linkages can be used to assess additional chemical classes, diseases, and/or geographic regions to characterize the relationship between chemicals, genetic variants, and disease outcomes in a population in order to protect vulnerable individuals and advance chemical risk and impact assessment in support of improving public health.

## Acknowledgements

This work was financially supported by the SPRINT project funded by the European Commission through Horizon 2020 (grant agreement no. 862568).

