## Supplementary material for "Genetic variability in pathways associates with pesticide-induced nervous system disease in the United States": SI Methods/Figures

### Contents

S1. Chemical-Gene data integration ... 2
S2. Pathway-Gene data integration ... 3
S2.1. Sensitivity analysis ... 3
S3. Chemical-Gene-Pathway-Disease integration ... 4
S3.1. Sensitivity analysis ... 5
S4. Overview of Pesticide-SNP-Disease linkages ... 7
S5. Spatialized Pesticide-SNP-Disease linkages ... 8
S5.1. SNP/Gene hits ... 9
S6. Comparisons between Pesticide-SNP-Disease linkages and disease occurrence by state 10
S7. State-to-state migration flows ... 18
S8. Random dataset generation ... 19
S9. LASSO regression ... 21
S10. High/Low probability state selection ... 23
S11. Frequent itemset mining analysis ... 23
S12. Priority list development ... 25
S12.1. SNP and gene list development ... 25
S12.2. Pesticide list development ... 26
S13. Data analysis and visualization ... 26
S14. References ... 27

### S1. Chemical-Gene data integration

Toxicology in the 21st Century (Tox21) is a federal research collaboration in the United States that screens large numbers of chemicals for activity, and the United States Environmental Protection Agency (US EPA) contributes in part to this initiative with an expanded assay set from the Toxicity Forecaster (ToxCast) initiative [1,2]. HTS data from Tox21/ToxCast version invitrodb v3.5 summary files (the most recent version at the time of analysis) were downloaded from the United States Environmental Protection Agency (https://www.epa.gov/chemical-research/exploring-toxcast-data-downloadable-data) and active chemical-gene associations were identified.

Chemical-assay hits were identified from the "EXPORT_LVL5&6" files by reducing chemical-assay associations to those with column "hitc" = 1, meaning the chemical was active in that assay. To avoid associations with potential cytotoxicity, only chemical-assay hits with an AC50 less than the AC50 for that chemical in cytotoxicity assays were kept (i.e. the AC50 for a given chemical-assay test was less than the AC50 for that same chemical in the "CytoPt.xlsx" file). Assays were reduced to those targeting human genes (based on data in the "gene_target_information_invitrodb_v3_5.xlsx" and "assay_annotation_information_invitrodb_v3_5.xlsx" files), and if an assay had multiple gene targets, these were counted as separate chemical-gene associations.

CTD is a publicly available database that curates information on chemicals, genes, and their linkages from the literature [3]. Chemical-gene interactions data were downloaded from the 28/10/2022 release of CTD (http://ctdbase.org/downloads). Data were reduced to species = Homo sapiens.

HTS and CTD chemical-gene associations were merged together based on CAS Registry Numbers (CASRN) and Entrez Gene IDs, following the approach outlined in Kosnik et al. 2019 [4]. The integrated HTS/CTD dataset was then reduced to pesticides used in the United States. These pesticides were identified based on data in the United States Geologic Survey (https://water.usgs.gov/nawqa/pnsp/usage), which provides the amount of pesticide applied in kilograms per county in the US between the years 1992 and 2019. At the time of analysis, data for 2019 was incomplete (50 total pesticides compared to 200+ for previous years), and was therefore excluded from the analysis. The USGS pesticides are given as compound names, so the

CompTox Chemicals Dashboard (https://comptox.epa.gov/dashboard/) was used to extract CASRNs for each pesticide based on the name.

### S2. Pathway-Gene data integration

Pathway-gene data from Reactome [5] and WikiPathways [6] were collected from the DAVID Knowledgebase on 16/11/2022 [7,8]. Pathway-Gene data from the Kyoto Encyclopedia of Genes and Genomes (KEGG) were collected using the pathfindR R package, version 1.6.3 [9].

Gene-disease data were collected from DisGeNET using the disgenet2r package version 0.99.3 [10] (https://www.disgenet.org/). These data include linkages curated from diverse sources, inferred from animal models or human phenotype associations, or identified through text-mining. These data were reduced to diseases in the Medical Subject Headings (MeSH) "Nervous System Disease" class (C10).

To identify pathways implicated in different nervous system diseases, ORA was used based on genes enriched in Pathway-Disease intersections. Enrichment was conducted using the disease_enrichment function in disgenet2r. All gene-disease associations were used because restricting to curated-only genes resulted in redundant Gene-Disease associations and fewer SNPs (as identified in subsequent steps). A sensitivity analysis was conducted to determine the best conditions to restrict Pathway-Disease associations to those with enrichment indicating the pathway is involved in a given disease without over-restricting associations to only those pathways and diseases with the largest numbers of starting gene associations.

#### S2.1. Sensitivity analysis

Enrichment was conducted with Pathway-Disease linkages identified with the FDR adjusted p-value for enrichment set at less than 0.05, 0.01, 0.001, or 0.0001. The additional requirement of a minimum of 3, 5, 10, 20, 35, or 50 genes overlapping between the pathway and disease was used as an additional parameter in the sensitivity analysis. A summary of these sensitivity analysis is in Figure S1. Based on these results, we determined that 3 or 5 genes in the Pathway-Disease intersection was too lenient a threshold and 35 or 50 genes at the intersection was too restrictive. We also determined that, with too restrictive a p-value, only Reactome pathways were left for each disease. Therefore, we considered Pathway-Disease linkages with an adjusted p-value of 0.01 or 0.001 and 10 or 20 genes overlapping in the subsequent Chemical-Pathway-Gene-Disease sensitivity analysis.

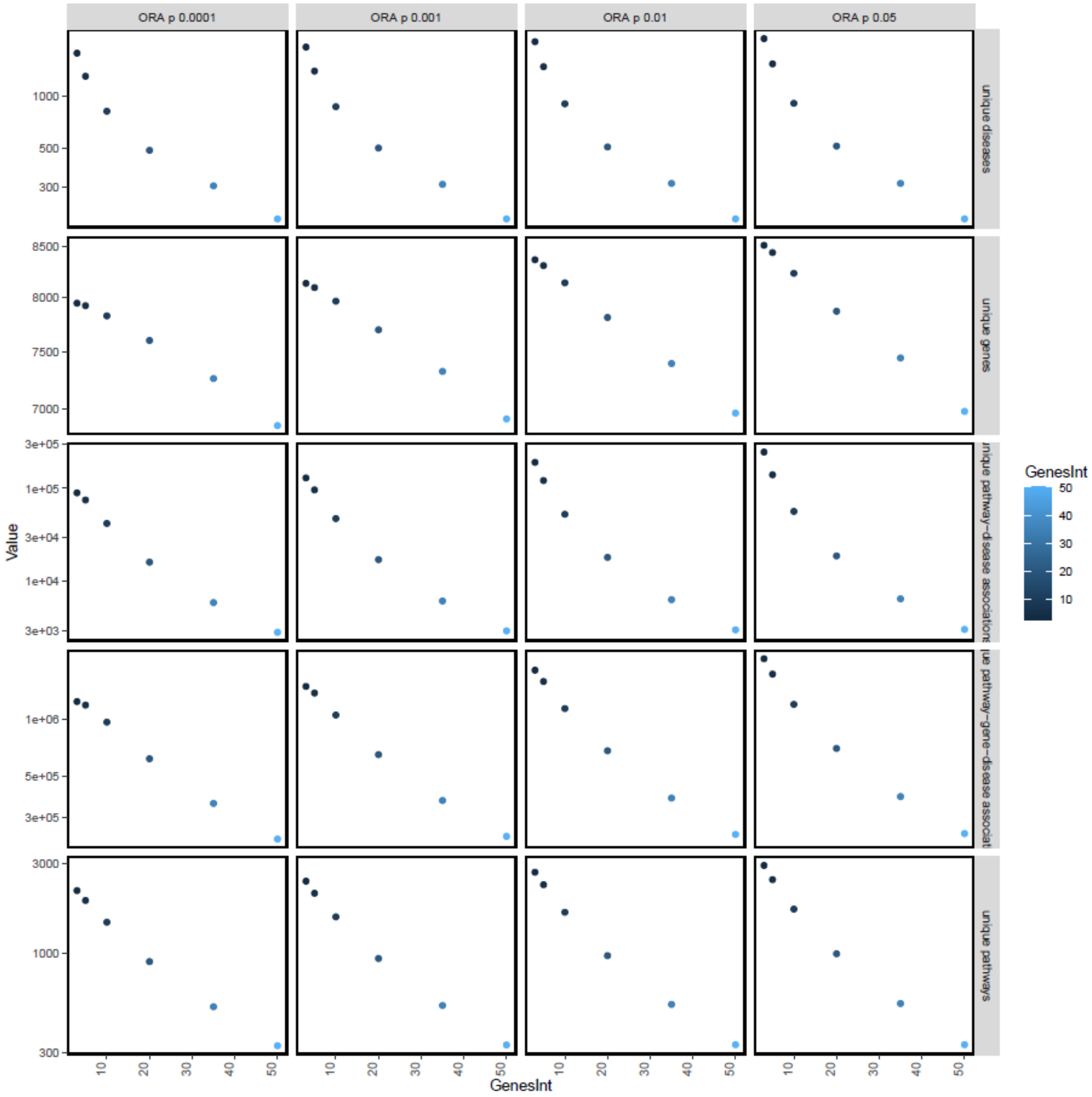


**Figure S 1.** Summary of attributes with different thresholds in the Pathway-Disease linkage. Columns = different p-value cutoffs in the over representation analysis. Rows = number of unique attributes in the dataset with each threshold requirement: unique Diseases, Genes, Pathway-Disease linkages, Pathway-Gene-Disease linkages, and Pathways following. GenesInt (Color/x-axis) = different requirements for the number of genes at the intersection of each Pathway-Disease linkage.

### S3. Chemical-Gene-Pathway-Disease integration

Fisher's exact test was used to determine significant overlap in the genes between Chemical-Gene linkages and Pathway-Gene-Disease linkages. Background was set as all disease-relevant genes in DisGeNET, and the fisher.test function in R was used. Significant linkages were identified according to the sensitivity analysis below.

#### S3.1. Sensitivity analysis

Using the Pathway-Disease linkages with an adjusted p-value of 0.01 or 0.001 and 10 or 20 genes overlapping as described in Section S2.1, Chemical-Pathway-Disease linkages were formed based on 3, 5, 10, or 15 genes overlapping between a chemical and a Pathway-Disease linkage and a Fisher's exact p-value of 0.05, 0.01, or 0.001. These results are in Figure S2.

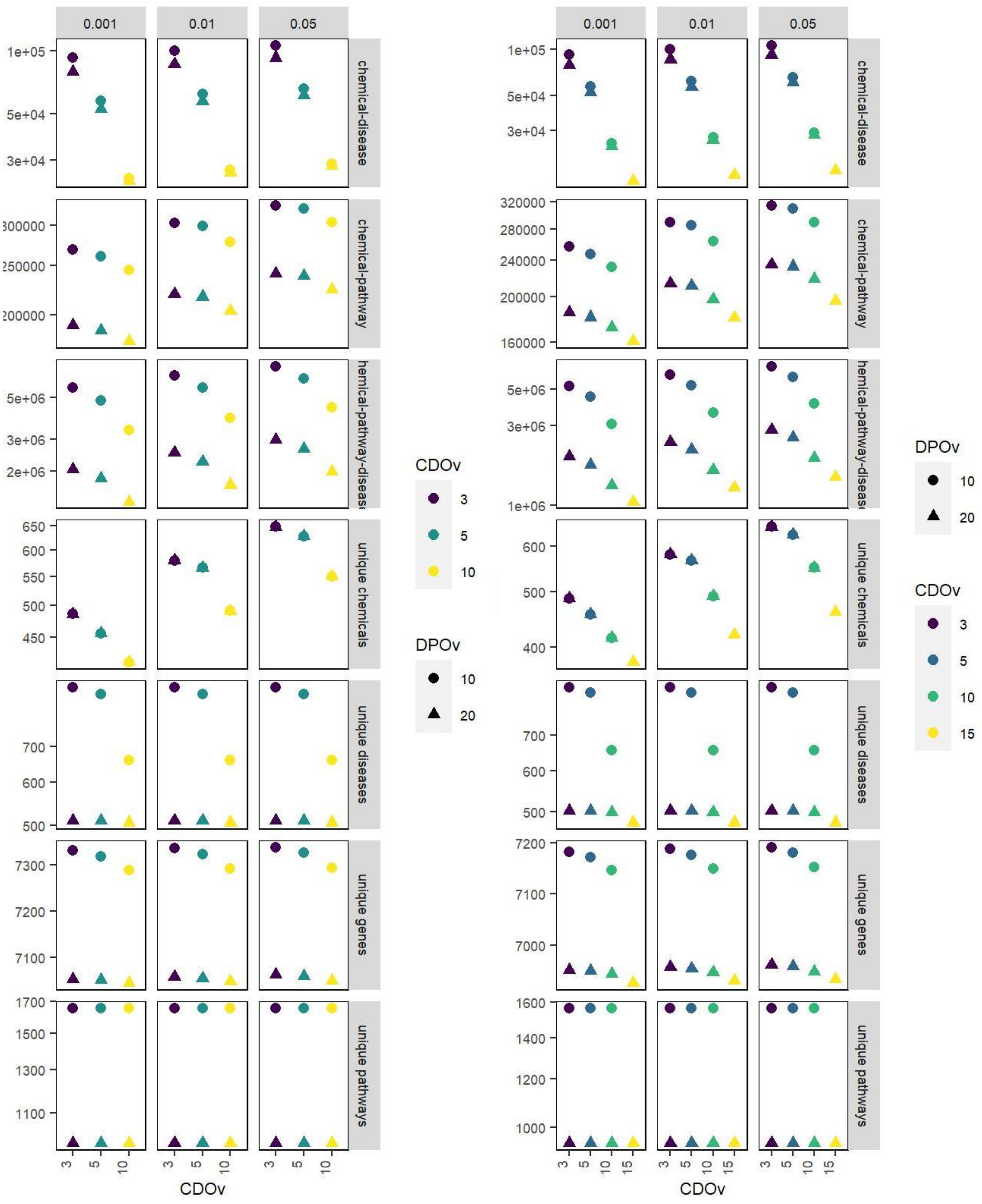


**Figure S 2.** Summary of Chemical-Pathway-Disease linkage attributes. A) Pathway-Disease ORA p-value requirement <0.01. Columns = different Fisher test p-value cutoffs in the Chemical-Disease linkage. Rows = number of unique attributes in the dataset with each threshold requirement. CDov (Color/x-axis) = different requirements for the number of genes at the intersection of each Chemical-

Disease linkage. DPOv (shape) = different requirements for the number of genes at the intersection of each Pathway-Disease linkage. B) Pathway-Disease ORA p-value requirement <0.001. Columns = different Fisher test p-value cutoffs in the Chemical-Disease linkage. Rows = number of unique attributes in the dataset with each threshold requirement. CDov (Color/x-axis) = different requirements for the number of genes at the intersection of each Chemical-Disease linkage. DPOv (shape) = different requirements for the number of genes at the intersection of each Pathway-Disease linkage.

Based on these results, we found that requiring >10 genes per Pathway-Disease overlap was considered too restrictive as it reduced the majority of pathway associations to Reactome pathways. Similarly, restricting the p-value to 0.001 removed many disease-relevant pathways (e.g., PPAR Signaling Pathway). Finally, requiring more than five genes per Chemical-Pathway-Disease linkage caused a large drop in the number of Chemicals and Chemical-Disease associations. Therefore, we set the Pathway-Disease associations with an ORA p-value < 0.001 and a requirement of >10 genes in the Pathway-Disease intersection, and the Chemical-Pathway-Disease associations with a Fisher's exact p-value < 0.01 and >5 genes in the Chemical-Pathway-Disease intersection.

### S4. Overview of Pesticide-SNP-Disease linkages

The final dataset includes 234 pesticides implicating 2304 total SNPs and 3244 total genes across 1230 toxicity pathways in six nervous system diseases of interest: Alzheimer's disease, Parkinson disease, multiple sclerosis, brain neoplasms, epilepsy, and migraine disorders (Figure 1, main text). These data form 1272 Pesticide-Disease associations with at least one SNP implicated and a median of 16 SNPs per Pesticide-Disease association (range 1-486) for a total of 36,691 unique Pesticide-SNP-Disease linkages. An additional 33,346 Pesticide-Gene-Disease linkages had no SNPs implicated. Pesticides with the most SNPs implicated include copper-based compounds (copper has 1017 SNPs implicated), natural pesticides like hydrogen peroxide and rotenone (1064 and 950 SNPs implicated, respectively), and synthetic pesticides like atrazine and paraquat (711 and 635 SNPs implicated, respectively). Alzheimer's disease, Parkinson's disease, and multiple sclerosis are the diseases with the most pesticides and SNPs implicated (233, 221, 229 pesticides, respectively, and 991, 409, and 644 SNPs). The full set of Pesticide-Pathway-Gene-SNP-Disease linkages is available in Table S2 (Excel file). Literature support for Pesticide-SNP-Disease linkages and/or Pesticide-Gene-Disease linkages is in Table S1.

**Table S1.** Literature support for Pesticide-SNP-Gene-Disease linkages

| Chemical | Gene | SNP | Disease | Reference |
|---|---|---|---|---|
| Pyrethroids | HLA-DRA | rs3129882 | Parkinson Disease | [11] |
| Organochlorines and/or organophosphates | ABCB1 | rs1045642, rs2032582 | Parkinson Disease | [12–14] |
| Organophosphates (Diazinon, Chlorpyrifos,) | PON1 | rs854560, rs662 | Parkinson Disease | [15,16] |
| Organophosphates (Chlorpyrifos) | NOS1 | | Parkinson Disease | [17] |
| Agricultural pesticides | ALDH2 | | Parkinson Disease | [18] |
| Paraquat | APEX1 | rs1130409 | Parkinson Disease | [19] |
| Maneb, Paraquat | NFE2L2 | rs6721961 | Parkinson Disease* | [20] |
| Paraquat | PPARGC1A | rs6821591, rs8192678 | Parkinson Disease** | [20] |
| Household pesticide products | MANBA | rs7665090 | Multiple Sclerosis*** | [21] |
| Copper | CYP2D6 | | Alzheimer's Disease | [22] |
| Copper | GSTP1 | rs1695, rs1138272 | Alzheimer's Disease | [22] |

* This SNP may delay Parkinson Disease onset (discounting pesticide exposures)
** Parkinson Disease-like symptoms, not the disease itself
*** Paediatric-onset multiple sclerosis

While there are limited data on SNP-Disease or Gene-Disease interactions in pesticide-exposed populations, we found good agreement between the linkages we formed and SNPs implicated in differential population susceptibility to pesticides in the literature (SI Table S1). The only SNPs we identified in the literature but not in our dataset were SNPs not incorporated in DisGeNET, the SNP database we used to form the linkages (e.g., rs671 in ALDH2 [18]; ALDH2 was implicated in our dataset with different SNPs). This validation of the linkages we formed increases confidence in our dataset.

### S5. Spatialized Pesticide-SNP-Disease linkages

Data on pesticide application between 1992 – 2018 for 3,065 counties in the contiguous US was available in USGS (https://water.usgs.gov/nawqa/pnsp/usage/maps/county-level/). Because disease data from Global Burden of Disease is only available per state, pesticide application data were aggregated across counties with the amount of pesticide applied per year per state as the metric for pesticide use. If the kilograms of pesticide applied was reported with both a high and low value estimate rather than a single value (about 20% of county-level data entries), the median was taken. This left a final dataset with the amount of pesticide applied per state per year for 1992 – 2018. Many pesticides were applied in all states in all years including 2,4-dichlorophenoxyacetic acid (2,4-D), dicamba, paraquat, chlorpyrifos, and glyphosate, which was also used at a greater mass than other pesticides, (4.5 times greater than the next most-applied pesticide, 2,4-D). All six nervous system diseases occurred in all states. Based on where pesticides were applied, SNP-disease associations were mapped to that state (e.g., Copper – rs669 – Alzheimer's disease was mapped to all states where copper was applied). The number of unique pesticides and kilograms/square mile of pesticide applied are in Figure S3, along with the total number of SNPs, and genes implicated and the total SNP and gene hits.

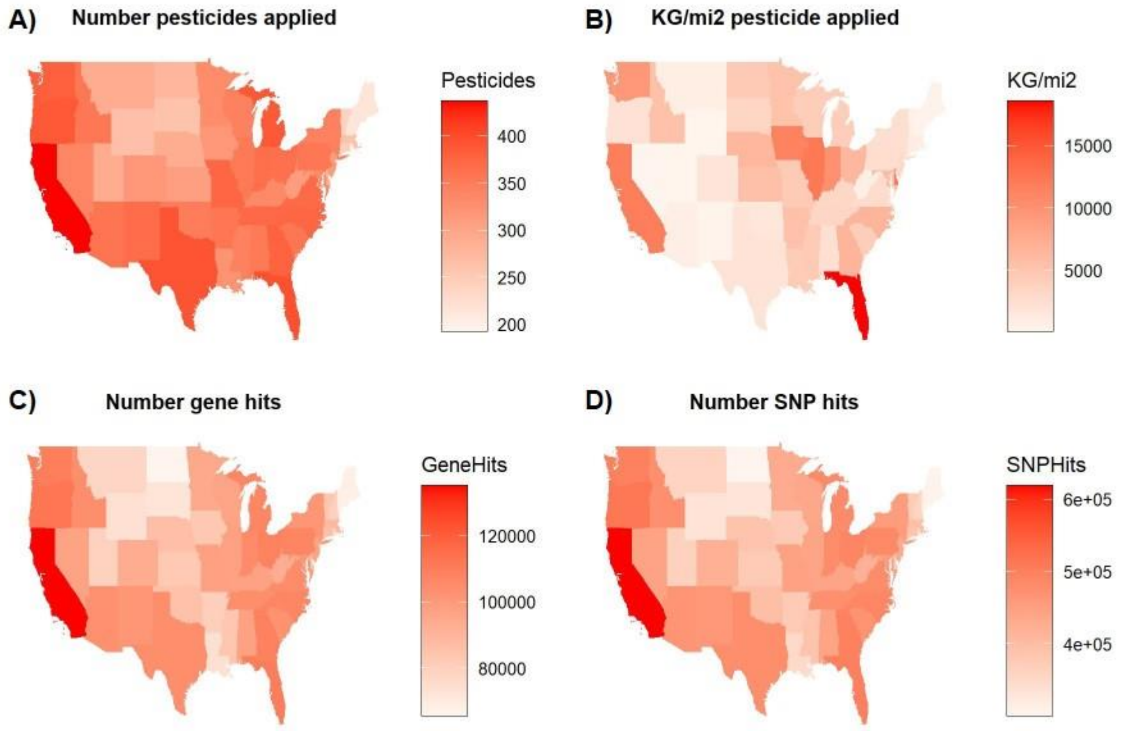


**Figure S 3** Spatialized pesticide, gene, and SNP data for pesticides applied in the US in 1992 – 2018. A) Number of different pesticides applied per state in 1992 – 2018. Note: these are all pesticides applied, not just pesticides with SNP-Disease associations. B) kilograms/square mile of pesticides applied in 1992 – 2018. C) Gene hits from pesticides applied in 1992 – 2018, D) SNP hits from pesticides applied in 1992 – 2018 (see Figure S4 for an explanation of SNP/gene hits).

#### S5.1. SNP/Gene hits

As multiple pesticides can implicate the same SNP/gene and the same pesticide can be applied in multiple years, the same SNP/gene can have multiple "hits" in a single year and across years. Therefore, to consider the cumulative potential for a SNP/gene to be affected by pesticide application, we assess SNP/gene frequency over years based on pesticide-induced hits. An example figure of SNP hit calculation is shown in Figure S4.

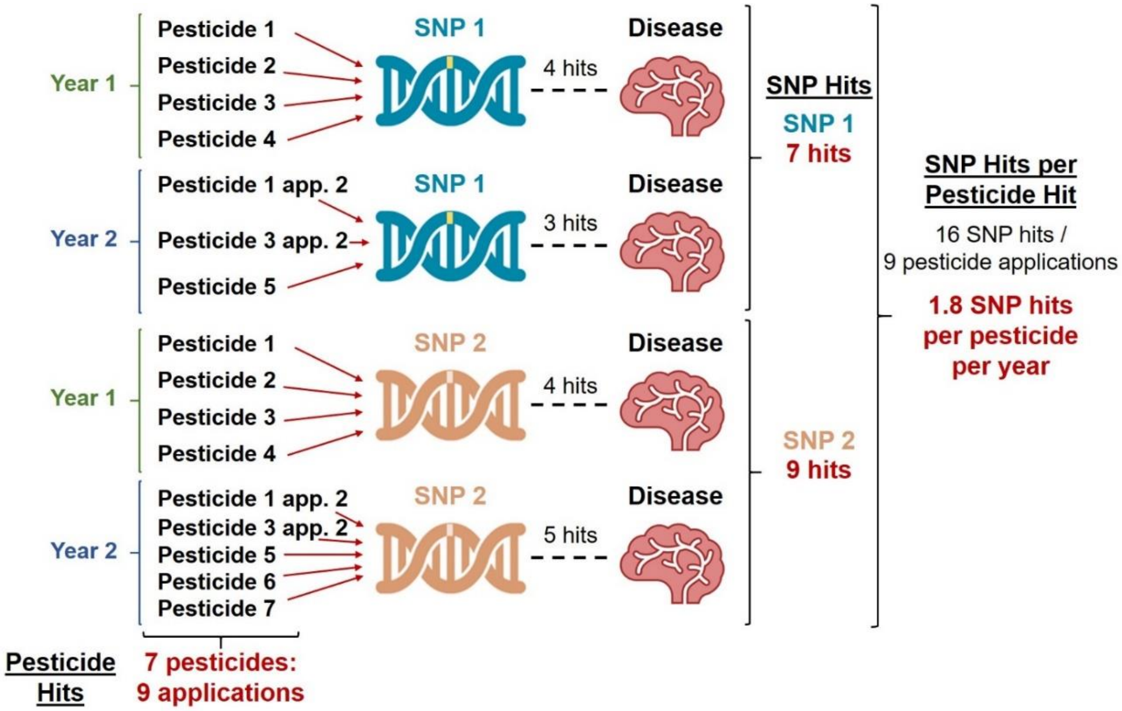


**Figure S 4.** Example diagram of SNP hit calculations. Each SNP can be affected by multiple pesticides in the same year (e.g., SNP 1 is affected by four pesticides in Year 1, hence has four hits) and over years (e.g., SNP 1 is affected by 3 pesticides in Year 2, hence three hits). Each time a pesticide is applied is a potential SNP hit, so each year of application is counted distinctly. Therefore the total hit frequency per SNP can be calculated (Total hits) or per pesticide per year (e.g., SNP hits/pesticide hits = 16 hits/9 times pesticides were applied = 1.8 SNP hits/pesticide-year).

### S6. Comparisons between Pesticide-SNP-Disease linkages and disease occurrence by state

For each year of disease data between 1992 and 2018, we compared the cumulative SNP hits from all pesticides applied prior to and including that year to the disease incidence and prevalence (i.e. for Alzheimer's disease incidence/prevalence in 2008, the SNPs and genes implicated based on pesticide use from 1992 – 2008 were compared). The Spearman's rho and robust linear model RMSE from each model is in Figure S 5. Generally, the relationship between the SNP hits/pesticide-year and disease occurrence got stronger over time. While the RMSE increased over time for Parkinson disease incidence and all disease prevalence, this is likely due to the increased range of disease cases between 1992 and 2018. For example, in 1992 the Parkinson disease incidence rate per 100000 people was between 11.2 – 21.2 compared to 31.6 – 68.5 for 2018. A similar trend was also observed for gene hits/pesticide-year and disease occurrence (Figure S 6).

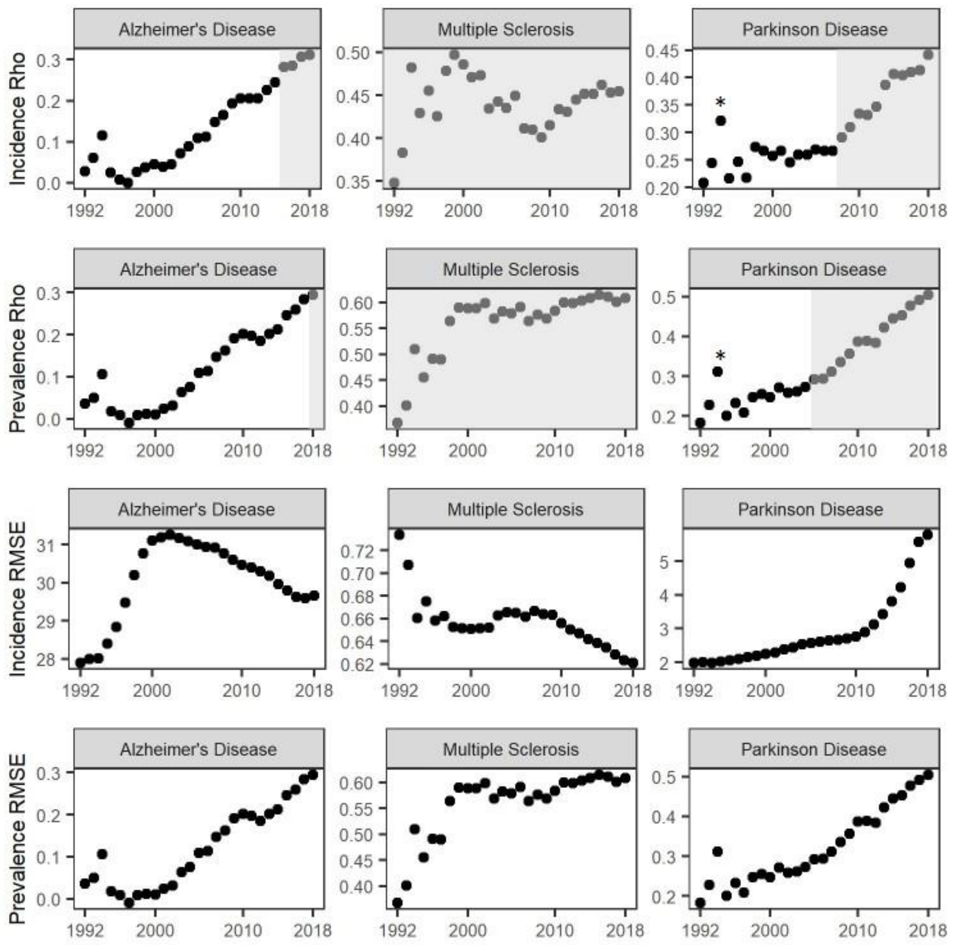


**Figure S 5.** Spearman's rho and robust linear model RMSE for SNP hits/pesticide-year vs disease incidence and prevalence over time. A grey box or asterisk indicate the spearman's correlation was significant for that model. The final year (2018) corresponds to the correlations in Figure S8.

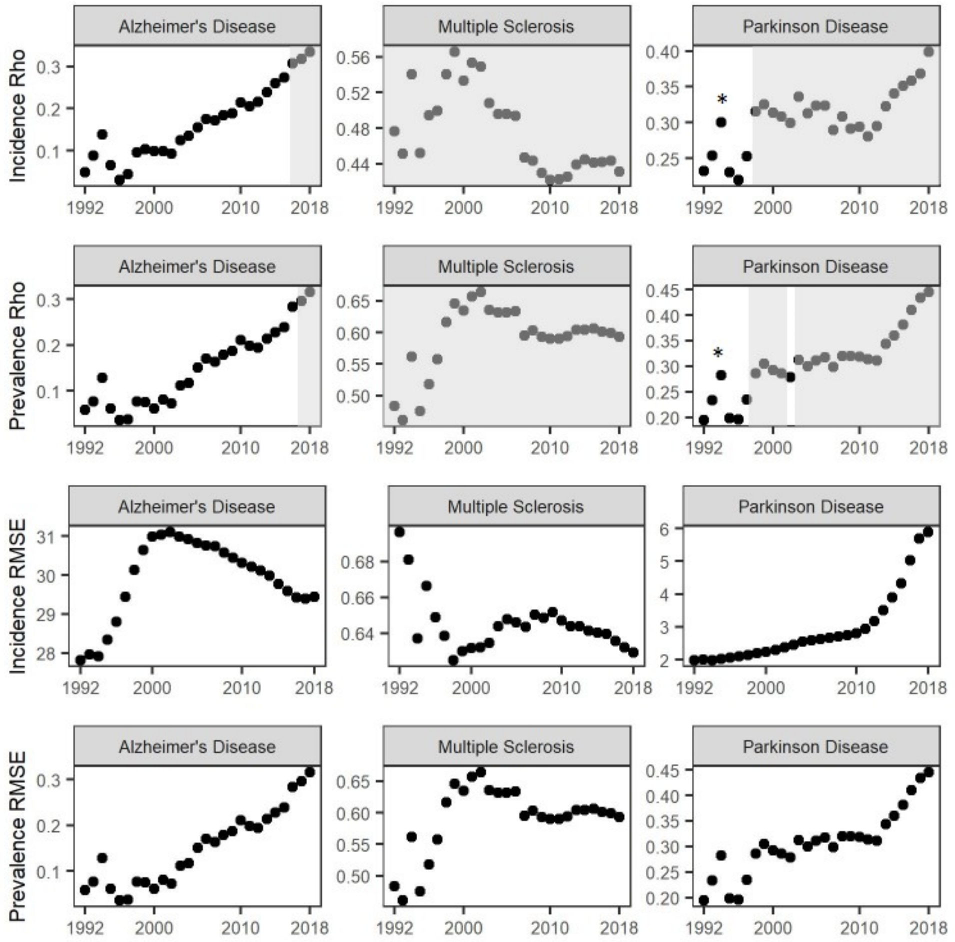


**Figure S 6.** Spearman’s rho and robust linear model RMSE for gene hits/pesticide-year vs disease incidence and prevalence over time. A grey box or asterisk indicate the spearman’s correlation was significant for that model. The final year (2018) corresponds to the correlations in Figure S9.

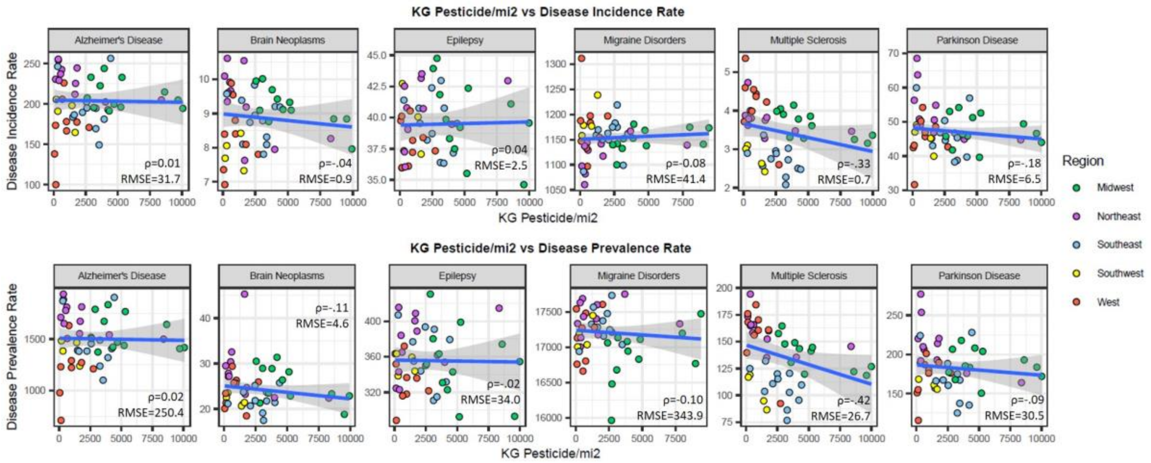


**Figure S 7.** Amount of pesticide (kg/m2) applied in 1992 – 2018 vs disease incidence or prevalence for the year 2018.

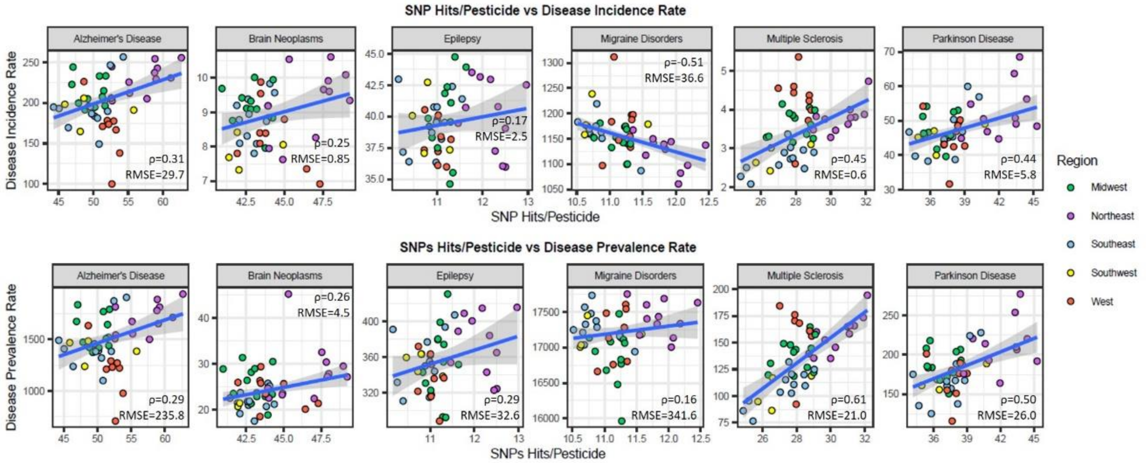


**Figure S 8.** Number of SNP hits/pesticide per year from pesticides applied in 1992 – 2018 vs disease incidence or prevalence for the year 2018.

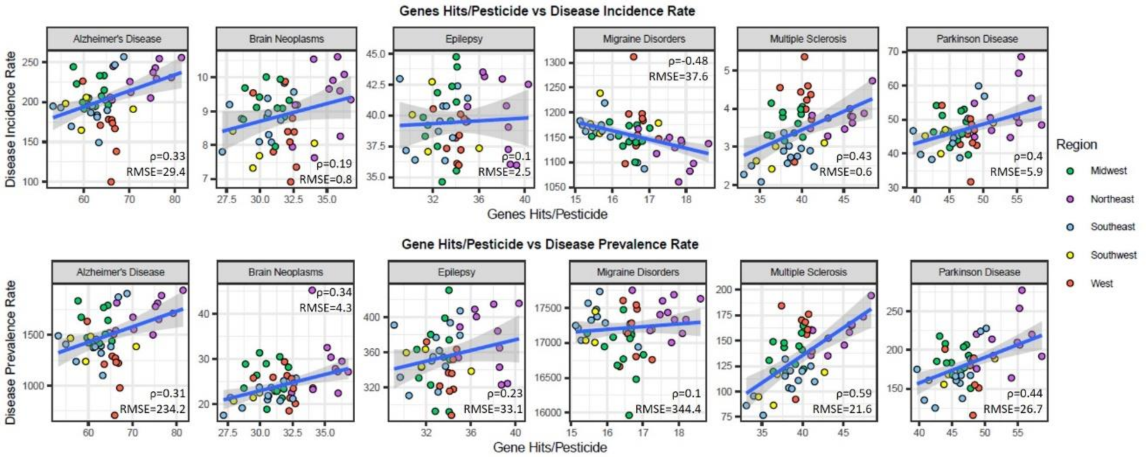


**Figure S 9.** Number of gene hits/pesticide per year from pesticides applied in 1992 – 2018 vs disease incidence or prevalence for 2018.

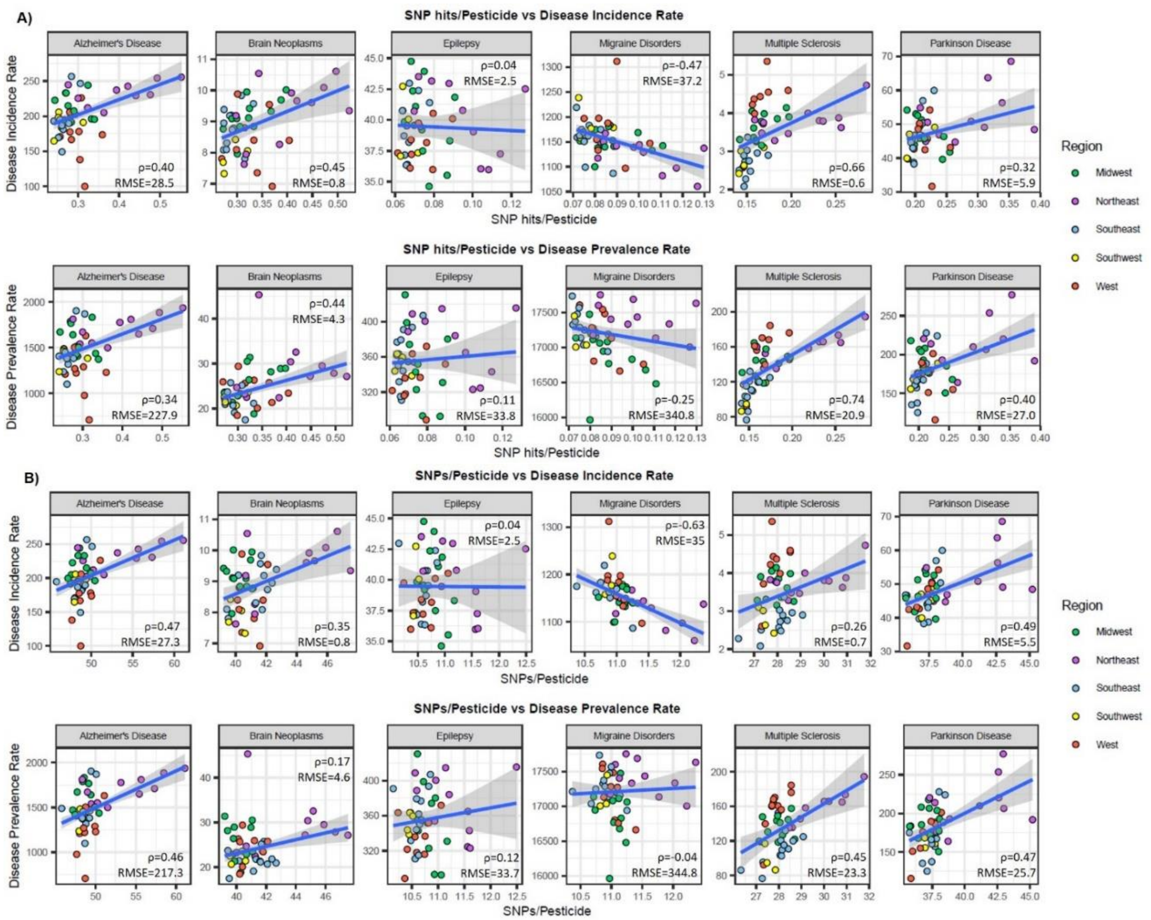


**Figure S 10.** SNPs/pesticide. A) The total number of SNP hits/pesticide across years 1992 – 2018 vs disease incidence or prevalence for 2018. B) The number of SNPs/pesticide applied (not SNP hits) in 1992 – 2018 vs disease incidence or prevalence for 2018.

We also compared each year of age-standardized disease incidence/prevalence and disease incidence/prevalence in age 55+ to the number of SNPs/pesticide-year (Figure S 11, Figure S 12). The general trend was the same as it was for the general population rate for each disease, but Alzheimer's disease was not significantly correlated. The relationship between disease incidence and prevalence and SNP hits/pesticide-year for 2018 is shown in Figure S 13. To better consider the potential that pesticide-SNP interactions increase the likelihood of developing nervous system disease in a general population (e.g., at younger ages as was found with multiple sclerosis SNPs, Table S1), we did not use age-standardized disease rates in our analysis.

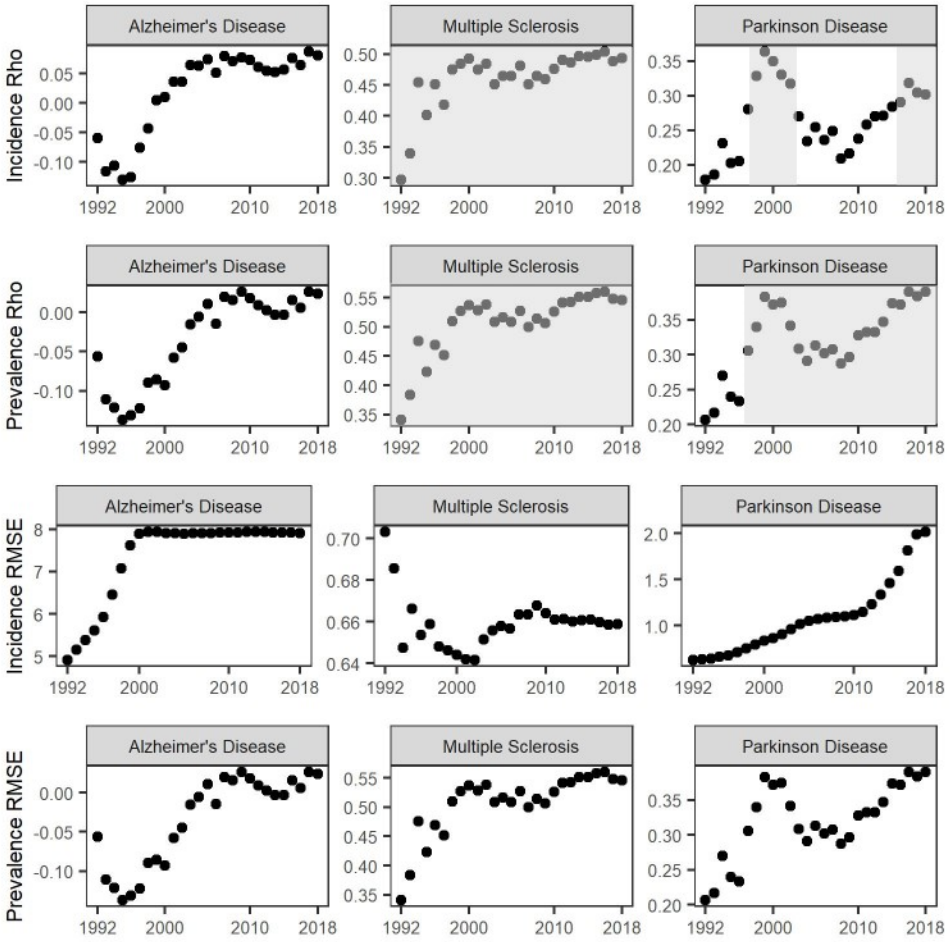


**Figure S 11.** Spearman's rho and robust linear model RMSE for SNP hits/pesticide-year vs age-standardized disease incidence and prevalence over time. A grey box or asterisk indicate the Spearman's correlation was significant for that model. The final year (2018) corresponds to the correlations in Figure S 13A.

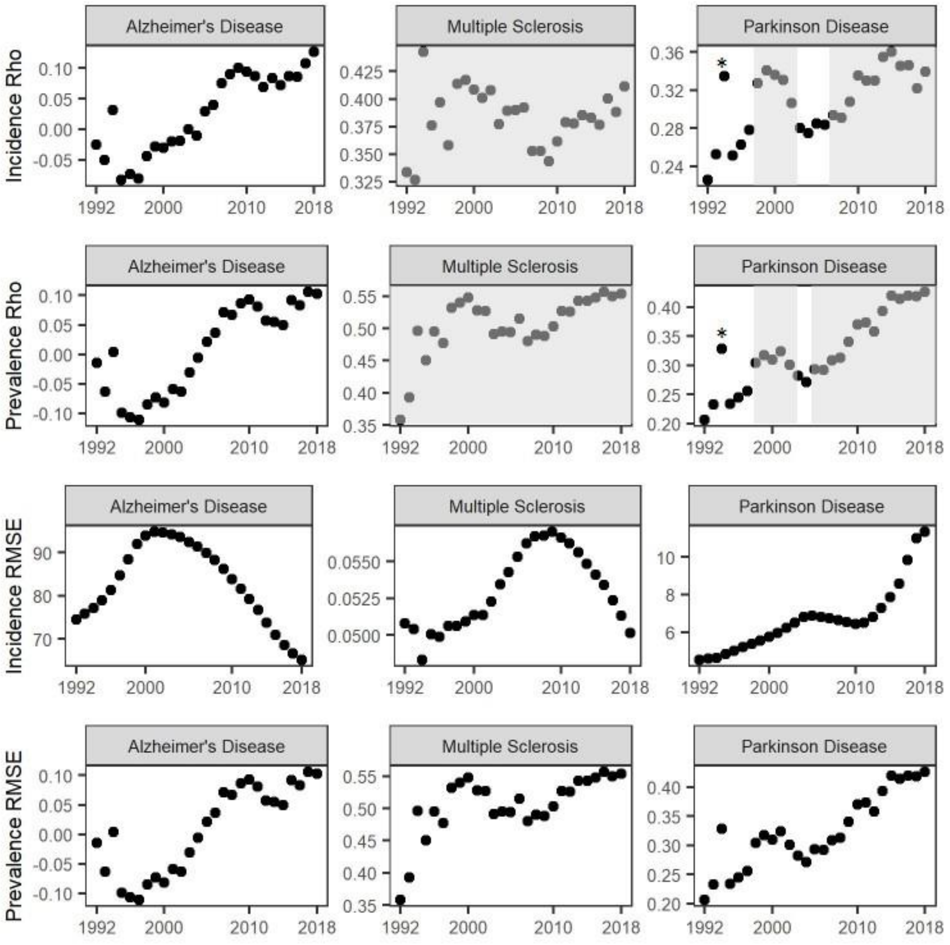


**Figure S 12.** Spearman's rho and robust linear model RMSE for SNP hits/pesticide-year vs age 55+ disease incidence and prevalence over time. A grey box or asterisk indicate the spearman's correlation was significant for that model. The final year (2018) corresponds to the correlations in Figure S13B spearman's correlation was significant for that model. The final year (2018) corresponds to the correlations in Figure S 13B.

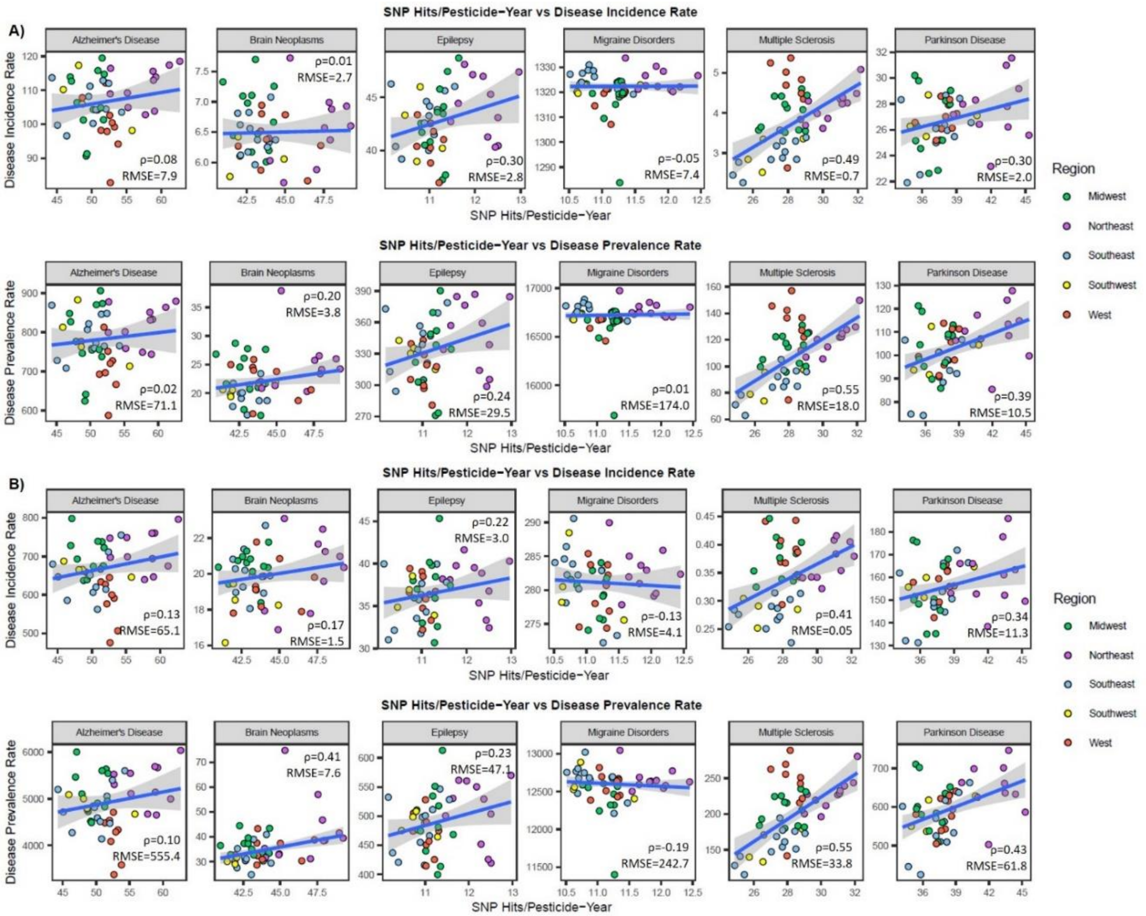


**Figure S 13.** A) The total number of SNP hits/pesticide-year for pesticides applied between 1992 – 2018 vs age-standardized disease incidence or prevalence for 2018. B) The total number of SNP hits/pesticide-year for pesticides applied between 1992 – 2018 vs age 55+ disease incidence or prevalence for 2018.

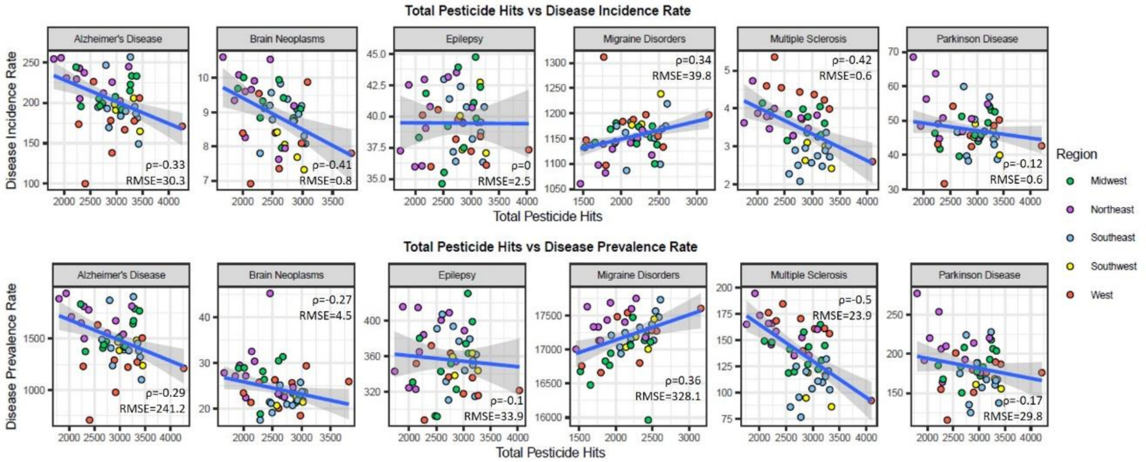


**Figure S 14.** The total number of times pesticides were applied in 1992 – 2018 vs disease incidence or prevalence for 2018.

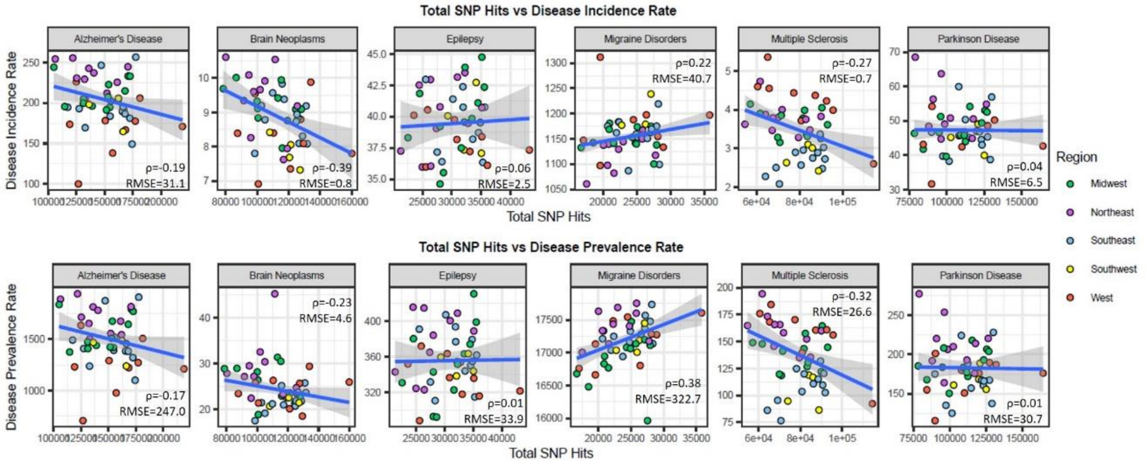


**Figure S 15.** The total number of SNP hits from pesticides applied in 1992 – 2018 vs disease incidence or prevalence for 2018.

The final dataset used for analysis associated all years of pesticide application data (1992 – 2018) to disease incidence and prevalence in 2018. For Alzheimer's disease, multiple sclerosis, and Parkinson disease, these associations were significant (Spearman p-value < 0.05) and the RLM equations were: 2.97x + 49.84 (Alzheimer's disease, incidence); 22.41x + 340.77 (Alzheimer's disease, prevalence); 0.22x – 2.77 (multiple sclerosis, incidence); 11.9x – 202.59 (multiple sclerosis, prevalence); 0.96x + 10.39 (Parkinson disease, incidence); 5.68x – 35.07 (Parkinson disease, prevalence).

### S7. State-to-state migration flows

One assumption of the analysis between SNP hits/pesticide-year versus disease incidence/prevalence is that the individuals in a state do not change over time. To test this assumption, we used census data on state-to-state migration flows for the US (https://www.census.gov/data/tables/time-series/demo/geographic-mobility/state-to-state-migration.html). These data provided information on how many people in a state lived in the same state the previous year, and were available for 2010 – 2018. Through this analysis, no more than 2.4% of the total US population had lived in a different state in the previous year (maximum 6.2% analyzing state-by-state) with most living in the same house as the previous year (84.5%).

Therefore, while there is likely some migration into and out of states for populations in this analysis, we expect it to account for a very small portion of the total population in a region.

**S8. Random dataset generation**

Based on the pesticides applied in each state in each year in 1992 – 2018, pesticides applied in each state were randomly shuffled (thus changing the SNPs implicated in each state), keeping the same number of pesticides applied overall in each state. For each randomly generated dataset, we compared the disease incidence/prevalence for 2018 with the number of SNP hits/pesticide-year. From this correlation, we checked the Spearman's correlation coefficient and developed a robust linear model, calculating the mean absolute error (MAE) and root mean square error (RMSE) for each random correlation. This was done 1000 times. The distribution of the random Spearman's rhos, MAEs, and RMSEs were compared to the same values for the true dataset. The distribution of RMSEs, MAEs, and spearman's rhos are in Figure S16 and S17. We found that the true Spearman's correlation coefficient was significantly higher ($p < 0.05$) than at least 75% of the randomly generated datasets for Alzheimer's disease and > 95% of the randomly generated datasets for multiple sclerosis and Parkinson disease. For 85% of the random datasets for Alzheimer's disease, 90% of the random datasets for Parkinson disease, and more than 99% of the random datasets for multiple sclerosis, the MAE and RMSE from the robust linear model were higher than for the true model values. For migraine disorders, brain neoplasms, and epilepsy the bootstrapping approach suggested that the relationships between pesticide usage and their incidence/prevalence could not be sufficiently explained by the Pesticide-SNP-Disease linkages on their own.

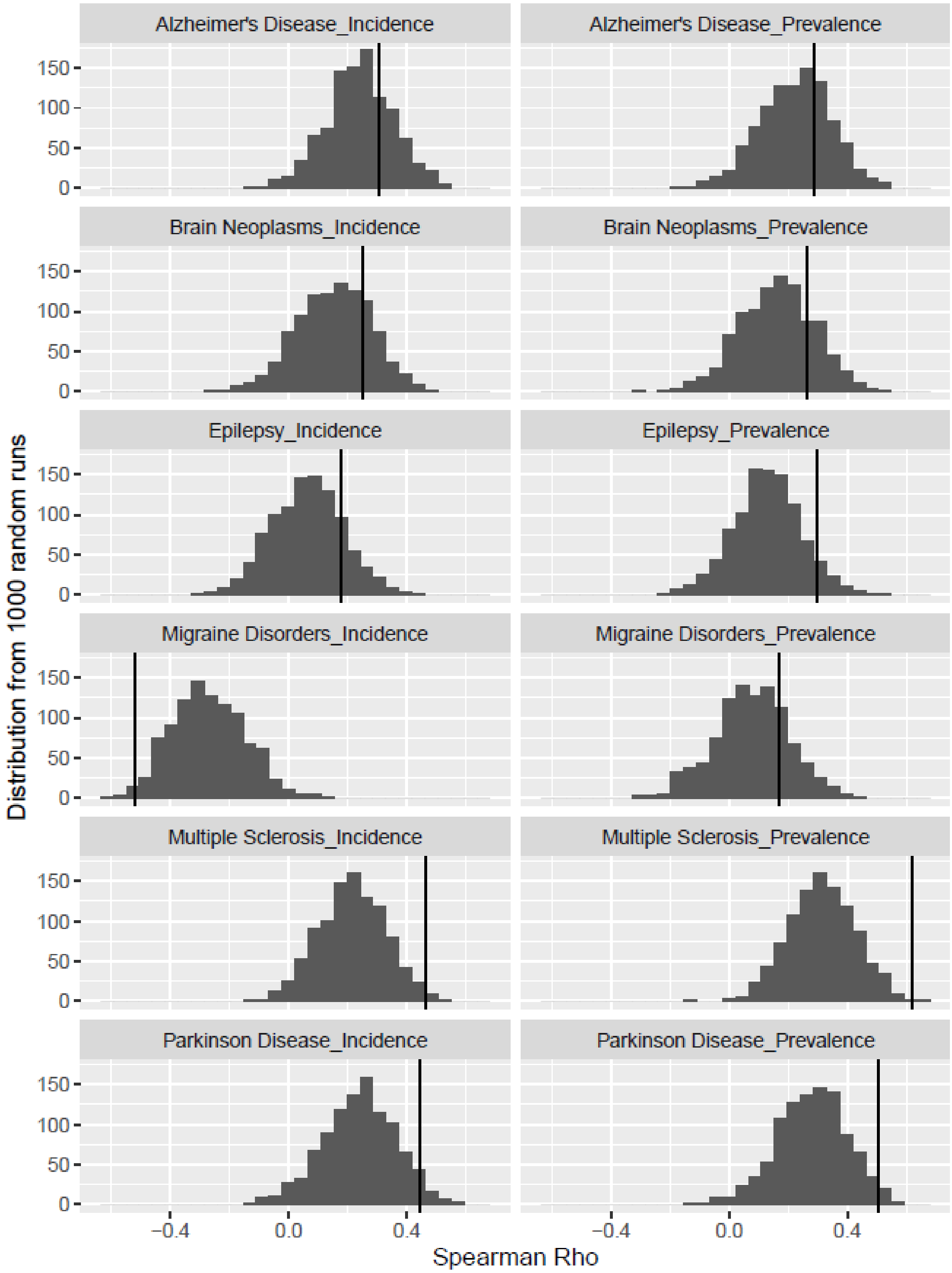


**Figure S 16.** Distribution of 1000 randomly generated dataset Spearman correlation coefficients versus the true value (black line).

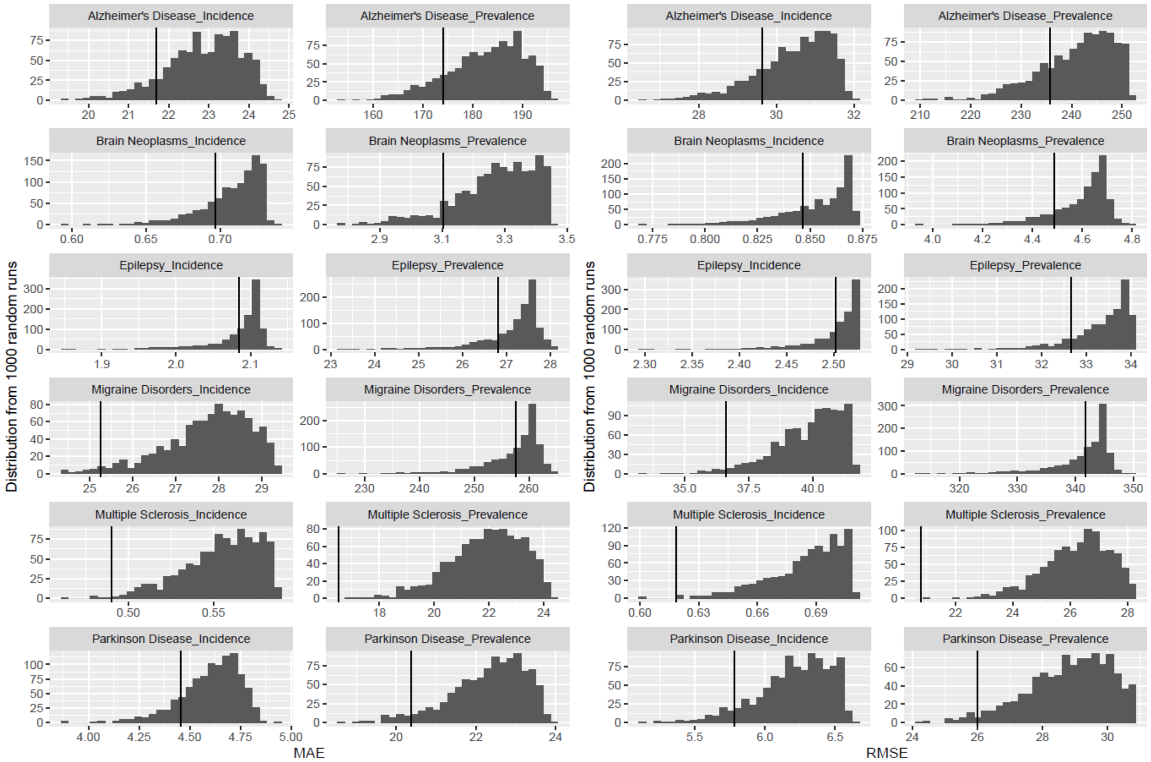


**Figure S 17.** Distribution of 1000 randomly generated dataset robust linear model mean absolute errors (MAE) and root mean squared errors (RMSE) versus the same, true values from the original dataset (black line).

### S9. LASSO regression

To assess the relationship between the different Pesticide-SNP hits and disease occurrence in each state, multi-linear regression models were built with the number of SNP hits implicated by each pesticide in each state as the predictor variable (i.e. the number of predictors was the number of pesticides). For all diseases, too many pesticides were implicated to build a normal multi-linear regression model owing to the limits on the degrees of freedom (48 states included in the model versus 216-227 predictors per disease).

LASSO regression was conducted using the c060 R package, version 0.2-9 (23) with the stabpath function. The size of the training set was 0.66 and 10000 steps were used. A separate model was built with the dependent variable as incidence or prevalence for Alzheimer's disease, multiple sclerosis, and Parkinson disease. The predictor variables were the number of SNP hits per pesticide per state (i.e. the number of predictors was the number of pesticides). In the final models, most pesticide predictors received coefficients. Alzheimer's disease incidence and prevalence had 218/227 pesticide predictors receive a coefficient while 205/216 multiple sclerosis pesticides and 212/223 Parkinson disease pesticides received a coefficient in the regularized LASSO regression for both disease incidence and prevalence. For all three diseases, the pesticides that did not receive a coefficient were used with the same application pattern in each state, meaning that there were no differences in the number of SNP hits per state (e.g., chlorpyrifos, glyphosate, paraquat, etc), or were pesticides only applied in one state (resmethrin, carbophenothion, dithiopyr).

By reducing the input for the robust linear model to just those compounds that received a coefficient in LASSO regression, we see a slight decrease in model performance, but models are still significantly correlated (spearman's correlation p < 0.05) and have better performance than most randomly-generated datasets (Figure S18)

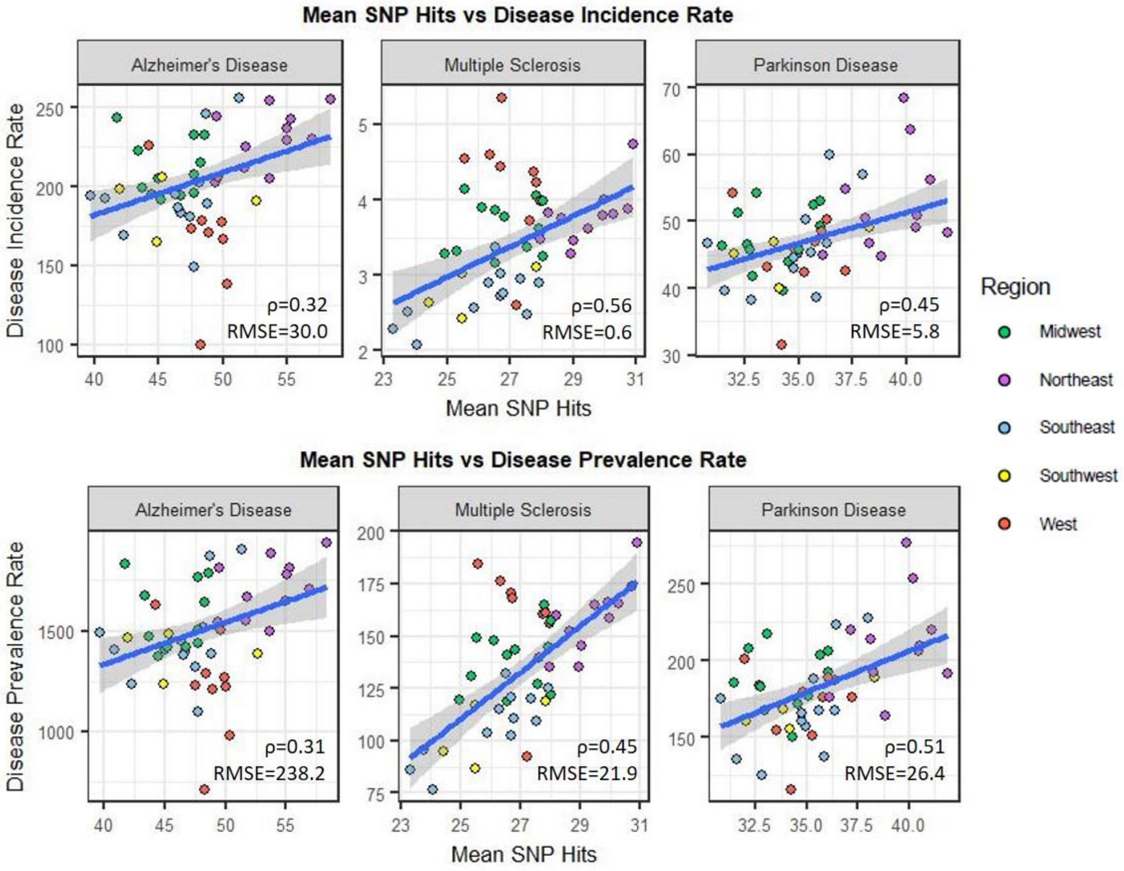


**Figure S 18.** SNP hits/pesticide-year for 1992 – 2018 with pesticide input reduced to non-zero coefficient pesticides from LASSO regression vs disease incidence or prevalence for the year 2018.

#### S10. High/Low probability state selection

To identify the high probability states (closer to the top right of the correlation plots) and low probability states for each disease (closer to the bottom left of the correlation plots), the values for the mean number of SNP hits per Pesticide-Disease linkage for each disease (x-axis in the correlation plots) were scaled between 0 and 1. The disease incidence and prevalence were also scaled between 0 and 1 (y-axis in the correlation plots), and the scaled incidence and prevalence were summed for each state to represent the disease occurrence. The disease occurrence value was then multiplied with the scaled SNP hits value to identify the high and low probability states for each disease (SI Figure S19).

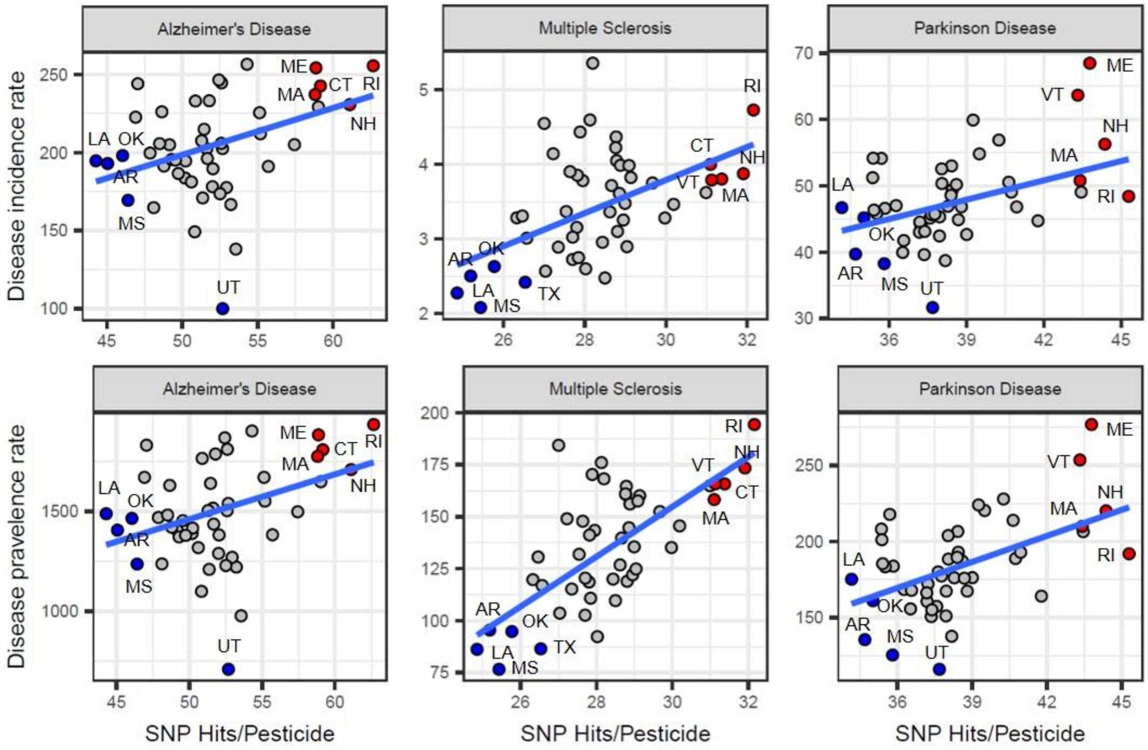


**Figure S 19.** High and low probability states for each disease. Red=High probability, blue = low probability. The red and blue states are the same for each column, but differ between diseases.

#### S11. Frequent itemset mining analysis

Association rules for pesticides were developed per state based on pesticides used in that state. For each state, each year of USGS data in 1992 – 2018 was used as an independent transaction with all unique pesticides used in that year as the input items (Figure S20). Rules between 2-4 pesticides long were identified with support set to 0.18 (meaning the combination had to appear in at least five years) and confidence set to 0.8. Analysis was done using the arules

R package version 1.7-5 [23] with the a priori function. Because we were more interested in the sets of pesticides identified than the directionality of the rule, only the resulting combinations of pesticides were kept, not the rules themselves (e.g., Rules A,B => C and B,C => A were considered the same rule: A,B,C).

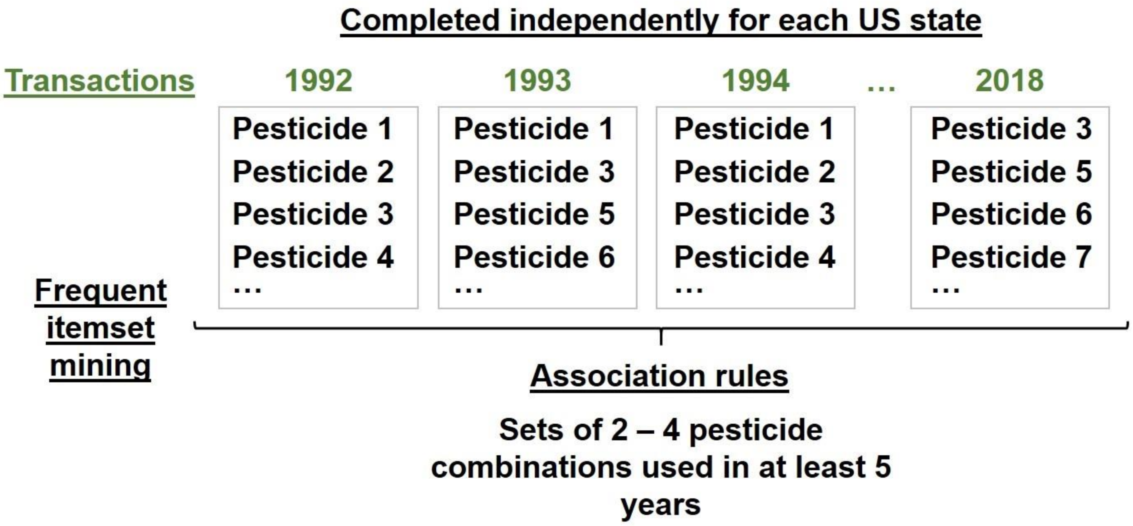


**Figure S 20.** Overview of the frequent itemset mining approach to identify frequently used pesticide combinations. Minimum support was set to 0.18 and confidence to 0.8.

The output of the frequent itemset mining analysis was a set of association rules per state describing recurring pesticide use patterns in 1992 – 2018. Association rules were mapped to diseases based on pesticides implicated in that disease (e.g., copper and fipronil were linked to Alzheimer's disease in our dataset, so the association rule copper – fipronil for Connecticut was linked to Alzheimer's disease). If an association rule for a state did not have any pesticides linked to that disease, then the association rule was excluded. If an association rule had at least one pesticide implicated in a disease (e.g., clofentezine – imidacloprid – vinclozolin was an association rule for Rhode Island and clofentezine and imidacloprid were linked to Alzheimer's disease in our dataset even though vinclozolin was not), then the association rule was kept because it is possible the currently unlinked pesticide could still be contributing to the disease, even if not directly linked in our dataset. Through this approach, association rules for high probability states could be compared to association rules for low probability states for each disease, and differences could be identified.

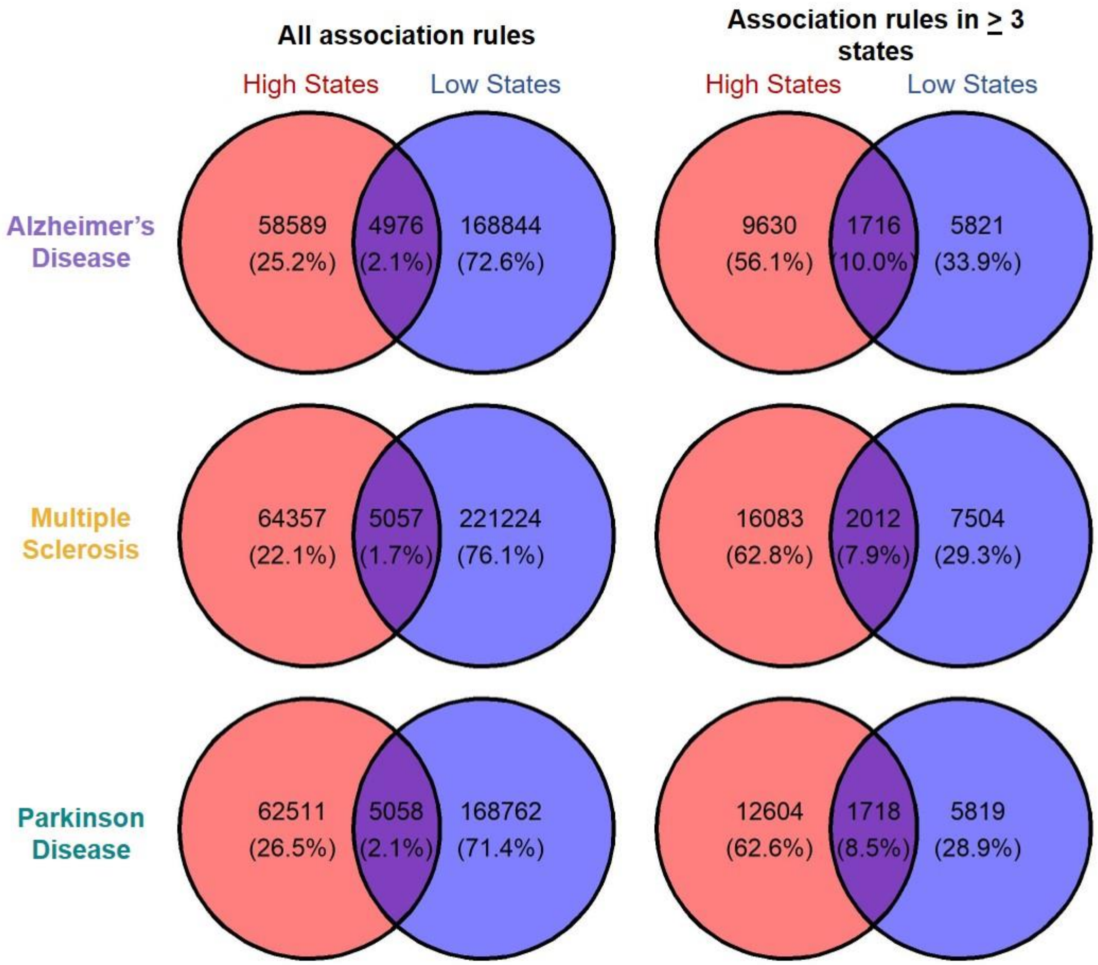


**Figure S 21.** Association rules per disease in high and low probability states or both. The left set of venn diagrams include all association rules implicated per disease in high and low probability states. The right set of venn diagrams only include association rules present in at least three high or three low probability states.

### S12. Priority list development

#### S12.1. SNP and gene list development

For each disease, a priority list of SNPs and genes was developed by counting the hit frequency: the number of hits (times a pesticide implicated that SNP/gene in that disease) across high probability states versus low probability states in 1992 – 2018. If the hit frequency for a SNP/gene in high probability states was higher than the hit frequency in low probability states, that SNP/gene was considered priority.

For each disease, the priority SNPs and genes were ranked based on the number of states implicating that SNP/gene and the number of pesticides present in association rules in high probability states. Rules were matched to SNPs/genes if at least one pesticide in that rule was linked to the SNP/gene in the Pesticide-Gene-SNP-Disease linkage. For each SNP/gene-rule combination, a rule value was assigned as follows:

1) If all pesticides in a rule implicated that SNP/gene in that disease in at least three high probability states, that SNP/gene-rule combination was assigned a rule value of five.

2) If fewer than all pesticides in a rule implicated that SNP/gene in that disease in at least three high probability states, that SNP/gene-rule combination was assigned a rule value of one.

Rules present in fewer than three states were not included in these calculations as they were not considered to distinguish differences in Pesticide-SNP/gene patterns in high and low probability states. Then, for each SNP/gene, the rank value was calculated as the sum of rule values multiplied by the hit frequency. Each disease priority list was then ordered by this rank value

#### S12.2. Pesticide list development

Priority pesticides were identified from pesticides acting on the high priority SNPs and genes identified in Section S12.1. Of these pesticides, the list was reduced to those pesticides implicated in three or more high probability states, occurring in association rules, and applied with greater mass in high probability states than low probability states. Then, pesticides were rank ordered with priority given to pesticides implicated only in association rules in high probability states but not low probability states, then the number of SNPs, genes, and difference in kilograms applied in high probability versus low probability states. Specific pesticide modes of action (MOAs) were determined from insecticide, fungicide, and herbicide resistance action committee classifications (IRAC, FRAC, and HRAC https://irac-online.org/mode-of-action/classification-online/, https://www.frac.info/fungicide-resistance-management/by-frac-mode-of-action-group, https://hracglobal.com/tools/hrac-mode-of-action-classification-2022-map).

### S13. Data analysis and visualization

Analyses were conducted using R version 4.2.2 [24]. Figures were generated using ggplot2, version 3.4.1 [25].
